# Satellite cell depletion in early adulthood attenuates muscular dystrophy pathogenesis

**DOI:** 10.1101/857433

**Authors:** Justin G. Boyer, Sarah Han, Vikram Prasad, Hadi Khalil, Ronald J. Vagnozzi, Jeffery D. Molkentin

## Abstract

Satellite cells are skeletal muscle resident stem cells that regenerate adult myofibers following an acute injury to muscle. Despite the assumption that the loss of satellite cells would be detrimental in a chronic regeneration-inducing muscle disease such as muscular dystrophy, this assumption has never been tested using mouse genetics. Here we generated a novel model of satellite cell ablation and crossed it with mouse models of muscular dystrophy to directly investigate how critical these cells are in maintaining muscle during a chronic degenerative disorder. Satellite cell deletion in 2-week-old young dystrophic mice provided noticeable improvements in histopathology and function, although at this early timepoint it was utimately detrimental because muscle size was not sufficient to permit survival. However, depletion of satellite cells beginning at 2 months of age in dystrophic mice provided similar histological and functional improvements but without compromising muscle size. The improved profile showed fewer damaged fibers, less myofiber central nucleation, increased sarcolemma integrity, decreased fibrosis and a dramatic size increase in the remaining myofibers. At the functional level, young adult dystrophic mice lacking satellite cells performed significantly better than those with satellite cells when exercised on a treadmill. Thus, loss of satellite cells during early adulthood in dystrophic mice produces an unexpected protective effect.

## Introduction

Muscular dystrophies (MD) are a collection of heterogenous neuromuscular diseases characterized by myofiber necrosis and dropout leading to a reduced lifespan. Duchenne muscular dystrophy (DMD) is an X-linked recessive myopathy caused by mutations in the *dystrophin* gene. This most common form of MD has an early childhood onset and affected individuals have a shorter life span (Shieh, 2013). Common pathological features of DMD include delayed motor milestones, proximal muscle weakness, hypertrophied calves and cardiomyopathy. The loss of the dystrophin protein in the *mdx* mouse, the most frequently used model of DMD, leads to a wave of myofiber degeneration/regeneration associated with elevated to creatine kinase beginning at 3 weeks of age (DiMario et al., 1991; Duddy et al., 2015).

Mutations in the *δ*-sarcoglycan (*Sgcd*) gene lead to limb-girdle muscular dystrophy (LGMD) 2F, which has a varying disease onset ranging from early childhood to adulthood. Individuals with this form of MD show progressive weakness and wasting of proximal muscles, elevated creatine kinase levels and survival is dependent on the level of cardiac and respiratory muscle pathology involvement (Blain and Straub, 2011). The absence of *Sgcd* in mice (*Sgcd*^*−/−*^) leads to similar but more severe muscle pathology compared to *mdx* mice. Some pathological indices include the presence of necrotic fibers, increased fibrosis and inflammatory cell infiltration (Hack et al., 2000). Currently, there is no effective treatment for either DMD or LGMDs.

One of the defining features of skeletal muscle lies in its capacity to regenerate following an injury event, which is due to muscle resident stem cells called satellite cells (Yin et al., 2013). Satellite cells are efficient at repairing muscle following an acute injury (Dumont et al., 2015). During the course of MD, while myofiber degeneration is continuously occurring, satellite cells are actively attempting to repair or replace damaged myofibers. However, satellite cells from dystrophic muscles give rise to myofibers harboring the same mutation leading to constant cycles of degeneration/regeneration. A therapeutic strategy currently being explored for DMD involves boosting the regenerative capacity of skeletal muscle by increasing satellite cell numbers or improving endogenous satellite cell function (Dumont and Rudnicki, 2016; Wang et al., 2019). How satellite cells influence the pathological outcome and progression of MD is not well understood. To gain insight into the role of satellite cells in MD, previous studies have irradiated skeletal muscle to deplete these cells to slow regeneration in dystrophic mice. Unexpectedly, irradiation delayed dystrophic disease onset (Pagel and Partridge, 1999) and it reduced muscle damage as determined by the increased sparing of peripherally nucleated fibers relative to the contralateral control leg (Granata et al., 1998). One study even noted that the dystrophic myofibers from the irradiated leg were hypertrophied (Wakeford et al., 1991). The observed effects on dystrophic muscles caused by irradiation occurred despite the fact that this method fails to deplete all satellite cells and can negatively affect multiple cell types within the irradiated leg (Granata et al., 1998; Scaramozza et al., 2019). More recently, Rossi and colleagues demonstrated that deletion of Nfix, a transcription factor responsible for the transcriptional switch from embryonic to fetal myogenesis, improved several disease pathological hallmarks in dystrophic mice (Rossi et al., 2017). These data suggest that delaying the regenerative response in MD could be protective for myofibers.

Despite these findings, the general assumption remains that the loss of satellite cells in a chronic muscle disease such as muscular dystrophy would be detrimental. This assumption however, has never been directly examined in vivo, such as in a mouse model where satellite cells can be genetically ablated at different time points during the disease. To assess the contribution of satellite cells to the pathogenesis of muscular dystrophy, we generated a highly efficient, biologically relevant mouse model of satellite cell ablation, which we crossed with 2 different mouse models of MD. We uncovered an intrinsic beneficial adaptive response in skeletal muscles during MD when satellite cells were deleted. This response was highlighted by a remarkable improvement in muscle histology and improved neuromuscular function.

## Results

### Erk1/2 are required for satellite cell viability upon activation

We have previously shown that deletion of p38 mitogen-activated protein kinase (MAPK) from mice with muscular dystrophy produced a protective effect to ongoing disease (Wissing et al., 2014), thus here we investigated the role that extracellular signal-regulated kinases1/2 (Erk1/2) play in regeneration and MD. MAPKs a highly conserved network of successively acting kinases, one branch of which culminates in the phosphorylation and activation of Erk1/2 (Plotnikov et al., 2011). Using isolated myofiber cultures, Erk1/2 signaling was shown to be associated with satellite cell function (Yablonka-Reuveni et al., 1999). Here we examined the role that Erk1/2 might play in satellite cell activity in vivo during injury and with chronic MD. Immunoblot analysis using lysate from regenerating tibialis anterior (TA) showed increased Erk1/2 expression on days 3 and 7 following cardiotoxin injury, with a leveling of expression by day 14 (Figure 1A). These results are consistent with the concept that Erk1/2 play a role early during myogenesis. We then generated satellite cell-specific *Mapk3 (Erk1); Mapk1 (Erk2)* double knockout mice to study the function of the Erk1/2 signaling pathway in muscle progenitors in vivo.

**Figure 1.**
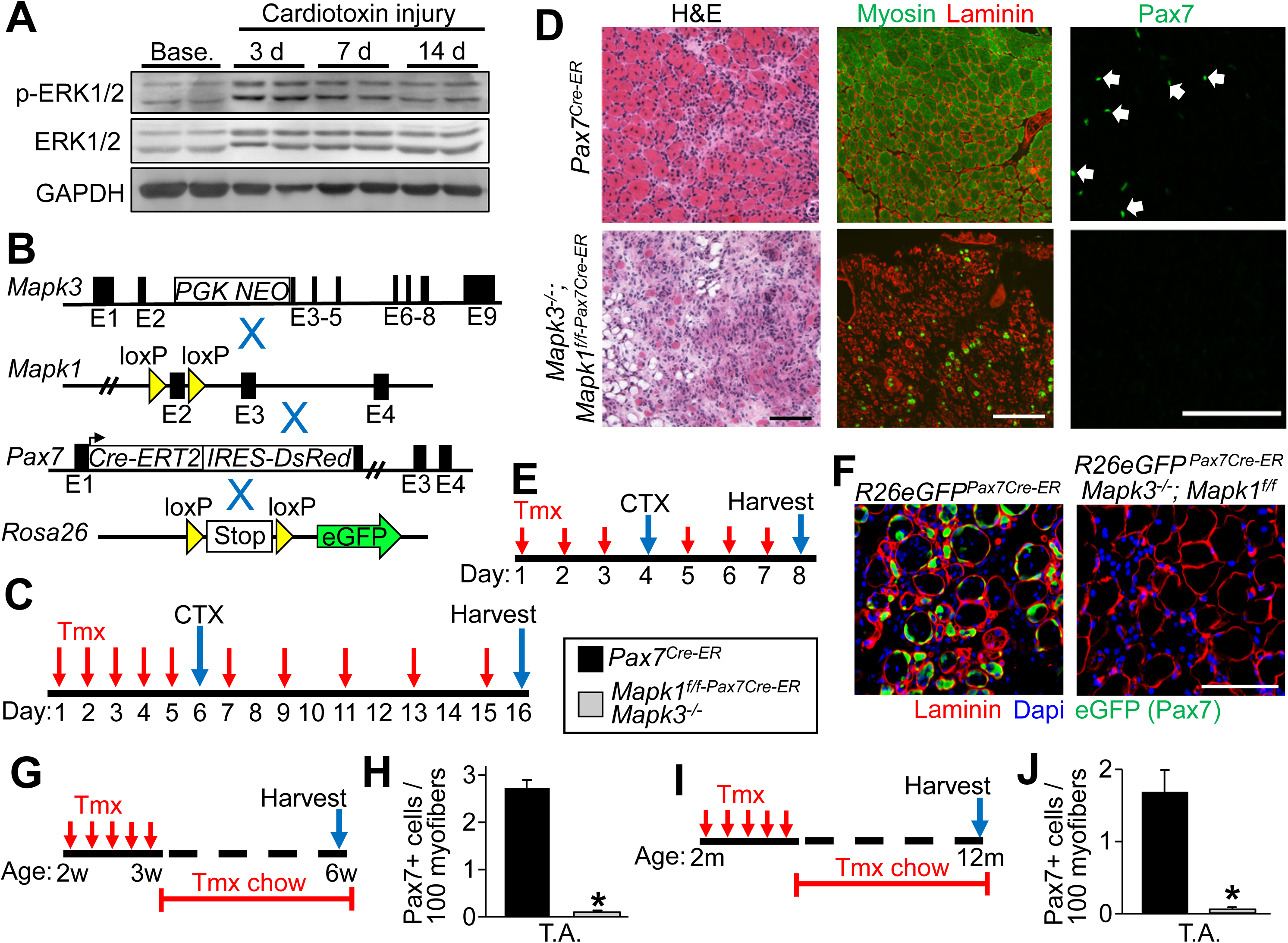
Erk1/2 are required for satellite cell viability upon activation. (A) Western blot analysis for the indicated proteins using tibialis anterior (TA) lysate from samples taken at baseline (base.) and following 3, 7 and 14 days (d) after cardiotoxin administration. Results from 2 different mice are shown. (B) Schematic representation of breeding of *Mapk3*^*−/−*^ and *Mapk1*-loxP(f)-targeted mice with the *Pax7*^*Cre-ERT2*^ mouse as well as with the Cre-dependent *R26eGFP* reporter mouse. (C) Schematic showing the timing of tamoxifen (tmx) and cardiotoxin (CTX) treatments in 2-month-old mice. (D) TA muscle sections of mice of the indicated genotypes stained with H&E (left panels), anti-myosin antibody (green) and anti-laminin antibody (red, middle panels) or with anti-Pax7 antibody (green, right panels) 10 days post CTX injury. Scale bars = 100 μm. (E) Experimental scheme showing that mice were injected with tmx for 3 days, then with CTX on day 4, followed by 2 more days of tmx injections. (F) Representative histological sections showing GFP fluorescence (green) and immunostained for laminin (red) 4 days post-CTX injection in mice of the indicated genotypes. Nuclei are stained with Dapi (blue). Scale bar = 100 μm. (G) Schematic representation of the tmx treatment regimen. Two week-old mice received a daily tmx injection for 5 consecutive days and were subsequently placed on tmx chow after weaning. (H) Quantification of satellite cells from uninjured TA muscle sections from mice of the indicated genotypes. n = 3 for both groups. Significance was determined using a Student’s t-test, *P < 0.05. (I) Schematic representation of tmx treatment. Two month-old mice received daily tmx injection for 5 consecutive days and were subsequently placed on tmx chow for 10 months. (J) Quantification of satellite cell numbers from uninjured TA muscle sections from 1-year old mice of the indicated genotypes. n = 3 for both groups. Significance was determined using a Student’s t-test, *P < 0.05. Data are plotted as the mean ± standard error of the mean (SEM) for all graphs.

*Mapk3*-null (*Mapk3*^−/−^) (Pages et al., 1999) mice were crossed with *Mapk1-loxP* site (fl) targeted mice (Samuels et al., 2008), which were also crossed with the tamoxifen inducible *Pax7*^*Cre-ERT2*^ mouse (Lepper et al., 2009) to allow for the deletion of *Mapk1* in satellite cells of adult mice following tamoxifen treatment (Figure 1B). Relative to controls, *Mapk1* transcripts were reduced 75% in freshly sorted satellite cells following 5 tamoxifen injections in *Mapk3*^−/−^; *Mapk1*^*f/f-Pax7Cre-ER*^ mice (Data not shown). To assess how the loss of Erk1/2 would impact satellite cell function, we treated 8 week-old *Mapk3*^−/−^; *Mapk1*^*f/f-Pax7Cre-ER*^ with tamoxifen and harvested muscles 10 days post-cardiotoxin injection (Figure 1C). Ten days after cardiotoxin injury, TA muscles from *Mapk3*^−/−^; *Mapk1*^*f/f-Pax7Cre-ER*^ mice were completely devoid of newly regenerating myofibers as demonstrated by H&E staining and by immunostaining for myosin heavy chain (Figure 1D). Immunohistochemistry for Pax7 expression was also absent in TA muscle 10 days after cardiotoxin injury in *Mapk3*^−/−^; *Mapk1*^*f/f-Pax7Cre-ER*^ mice. These results indicate that Erk1/2 are required in satellite cells for muscle regeneration.

*Mapk3*^−/−^; *Mapk1*^*f/f-Pax7Cre-ER*^ mice were also crossed with the *Rosa26 eGFP* (Yamamoto et al., 2009) Cre-dependent reporter mouse line to directly track satellite cells in vivo (Figure 1B). Muscle regeneration was assessed in 8 week-old *R26eGFP*^*Pax7Cre-ER*^; *Mapk3*^*−/−*^; *Mapk1*^*f/f*^ mice in which tamoxifen was first given to delete *Mapk1*, then cardiotoxin was given and the TA muscle was harvested 4 days later (Figure 1E). *R26eGFP*^*Pax7Cre-ER*^; *Mapk3*^*−/−*^; *Mapk1*^*f/f*^ injured muscles were indistinguishable from controls when comparing H&E stained histological sections 4 days post cardiotoxin injury (Data not shown). However, immunohistochemistry analysis revealed that with Mapk1/3 deletion no Pax7^+^ eGFP signal was observed surrounding necrotic fibers in *R26eGFP*^*Pax7Cre-ER*^; *Mapk3*^*−/−*^; *Mapk1*^*f/f*^ mice, but abundant Pax7^+^ myofibers were observed in control mice (Figure 1F). These data suggest that Erk1/2-deficient satellite cells die and are cleared following acute injury when activated.

To assess the need for Erk1/2 in satellite cells in uninjured muscles, *Mapk3*^−/−^; *Mapk1*^*f/f-Pax7Cre-ER*^ and control mice were treated with tamoxifen beginning at 2 weeks of age, a time when satellite cells are actively contributing to muscle growth (Pawlikowski et al., 2015; White et al., 2010). Tamoxifen treatment resulted in a 96% decrease in satellite cells in 6 week-old *Mapk3*^−/−^; *Mapk1*^*f/f-Pax7Cre-ER*^ mice (Figure 1G and H). Long-term analyses showed the complete depletion (97% decrease) of the satellite cell pool in 12-month-old *Mapk3*^−/−^; *Mapk1*^*f/f-Pax7Cre-ER*^ mice treated with tamoxifen starting at 2 months of age (Figure 1I and J). Singular deletion of *Mapk3* or *Mapk1* (using the LoxP-Cre strategy) did not affect satellite cell viability suggesting that these 2 kinases are redundant and that both genes must be deleted to lose satellite cells (Data not shown).

We crossed *Mapk3*^−/−^; *Mapk1*^*f/f*^ mice with a mouse containing a myofiber-specific Cre transgene in which the human skeletal α-actin promoter drives expression of the tamoxifen-inducible MerCreMer cDNA (Ska-MCM), which will show the consequences of depleting Erk1/2 in the late stages of myogenesis (Supplemental Figure 1B). We subjected *Mapk3*^−/−^; *Mapk1*^*f/f-Ska-MCM*^ mice to cardiotoxin injury and assessed the regenerative capacity of these mice as well (Supplemental Figure 1A). Histopathological analyses and quantification revealed that deletion of Erk1/2 in the later stages of myogenesis did not impact the regenerative response in *Mapk3*^−/−^; *Mapk1*^*f/f-Ska-MCM*^ mice relative to controls (Supplemental Figure 1C-F).

Satellite cells are active during the post-natal growth period and in adult muscle in response to injury. In mice satellite cell activity is minimal after 3 months of age where they likely remain in a more quiescent state (Pawlikowski et al., 2015). Therefore, we treated 6-to 8-month-old *Mapk3*^−/−^; *Mapk1*^*f/f-Pax7Cre-ER*^ and control mice with tamoxifen for 4 weeks. Our analysis revealed similar satellite cell numbers in the TA muscle sections of *Mapk3*^−/−^; *Mapk1*^*f/f-Pax7Cre-ER*^ and control mice at baseline without activation of these cells (Supplemental Figure 2A-C). However, 7 days post cardiotoxin injury these same *Mapk3*^−/−^; *Mapk1*^*f/f-Pax7Cre-ER*^ mice at this older age were devoid of any regenerating myofibers clearly demonstrating the need for Erk1/2 in satellite cell viability upon activation at essentially any age (Supplemental Figure 2D).

### Loss of satellite cells prior to disease onset reduces muscle damage in a mouse model of LGMD2F

Having established a novel model of satellite cell ablation, we used *Mapk3*^−/−^; *Mapk1*^*f/f-Pax7Cre-ER*^ mice to investigate satellite cell function in MD. We crossed *Mapk3*^−/−^; *Mapk1*^*f/f-Pax7Cre-ER*^ mice onto the *Sgcd*^*−/−*^ background (Figure 2A, (Hack et al., 2000)) and depleted satellite cells at various disease stages. Similar to *mdx* mice, *Sgcd*^*−/−*^ mice do not show pathological features of disease at 2 weeks of age (Data not shown). Beginning at 3 weeks however, an initial phase of myofiber degeneration occurs leading to a robust regenerative response highlighted by de novo myofiber replacement and hyperplasia.

**Figure 2.**
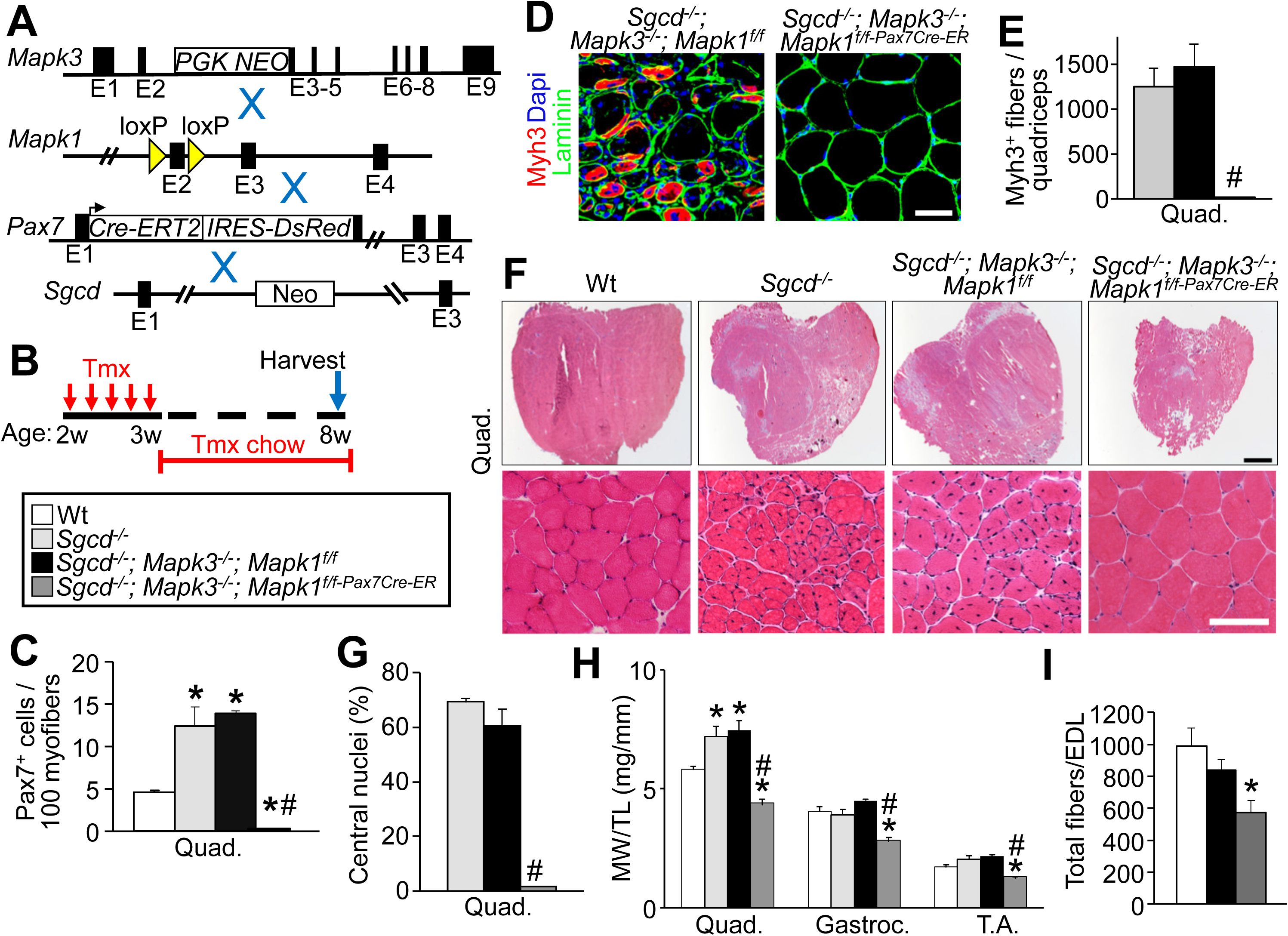
Satellite cell depletion prior to myofiber degeneration in *Sgcd*^*−/−*^ mice. (A) Schematic of breeding tmx-inducible *Pax7*^*Cre-ERT2*^ mice with *Mapk3*^*−/−*^ and *Mapk1*-loxP(f)-targeted mice. These lines were crossed onto the *δ-sarcoglycan*–null (*Sgcd*^*−/−*^) background. (B) Schematic of the tmx treatment. Two week-old mice received a daily tmx injection for 5 consecutive days and were subsequently placed on tmx chow after weaning. (C) Quantification of Pax7 positive satellite cells in muscle sections from the quadriceps (quad) of mice with the indicated genotypes. n = 3 for *Sgcd*^*−/−*^ samples and n = 4 for all other genotypes. One-way ANOVA with Tukey’s multiple comparisons test was used to determine significance, *P < 0.05 versus wild type (wt) controls, # P < 0.05 versus disease controls. (D) Representative quad muscle sections immunostained for Myh3 (red) and laminin (green) in 2 month-old mice of the indicated genotypes. Dapi stained nuclei are in blue. Scale bar = 50 μm. (E) Quantification of the number of Myh3 positive fibers in the quad muscle sections of mice with the indicated genotypes. n = 3 for *Sgcd*^*−/−*^ and *Sgcd*^*−/−*^; *Mapk3*^−/−^; *Mapk1*^*f/f*^ samples and n = 4 *Sgcd*^−/−^; *Mapk3*^−/−^; *Mapk1*^*f/f-Pax7Cre-ER*^. One-way ANOVA with Tukey’s multiple comparisons test was used to determine significance, # P < 0.05 versus disease controls. (F) Representative H&E stained histological sections of the quad muscle from mice of the indicated genotypes. Scale bars = 100 μm. (G) Quantification of the number of myofibers with centrally located nuclei in quad muscle sections from mice of the indicated genotypes. n = 4, *Sgcd*^*−/−*^; n = 3, *Sgcd*^*−/−*^; *Mapk3*^−/−^; *Mapk1^f/f^*; n = 5, *Sgcd*^*−/−*^; *Mapk3*^−/−^; *Mapk1*^*f/f-Pax7Cre-ER*^. One-way ANOVA with Tukey’s multiple comparisons test was used to determine significance, # P < 0.05 versus disease controls. (H) Muscle weights (M.W.) normalized to tibia length (T.L.) from mice of the indicated genotypes at 2 months of age. n = 5, wt; n = 5, *Sgcd*^*−/−*^; n = 3, *Sgcd*^*−/−*^; *Mapk3*^−/−^; *Mapk1^f/f^*; n = 7, *Sgcd*^*−/−*^; *Mapk3*^−/−^; *Mapk1*^*f/f-Pax7Cre-ER*^. A one-way ANOVA with Tukey’s multiple comparisons test was used to determine significance, *P < 0.05 versus wt, # P < 0.05 versus disease controls. (I) Quantification of the total myofibers present in the extensor digitorum longus (EDL) muscle from 2-month-old mice of the indicated genotypes. n = 3, wt; n = 4 *Sgcd*^*−/−*^; *Mapk3*^−/−^; *Mapk1^f/f^*; n = 4, *Sgcd*^*−/−*^; *Mapk3*^−/−^; *Mapk1*^*f/f-Pax7Cre-ER*^. A one-way ANOVA with Tukey’s multiple comparisons test was used to determine significance, *P < 0.05 versus wt. Data represent mean ± SEM for all graphs.

Here we first treated control and *Sgcd*^*−/−*^; *Mapk3*^−/−^; *Mapk1*^*f/f-Pax7Cre-ER*^ mice with tamoxifen at 2 weeks of age followed by tissue harvest 6 weeks later at what is typically peak disease (Figure 2B). The three control groups in our study were as follows: wild type healthy controls, *Sgcd*^*−/−*^ mice and *Sgcd*^*−/−*^; *Mapk3*^−/−^; *Mapk1*^*f/f*^ mice that were Cre negative. Throughout the study, no overt differences between *Sgcd*^*−/−*^ and *Sgcd*^*−/−*^; *Mapk3*^−/−^; *Mapk1*^*f/f*^ dystrophic mice were observed. We did not believe it necessary to include *Sgcd*^*−/−*^ mice harboring the *Pax7*^*Cre-ER*^ allele alone (Lepper et al., 2009) given that our objective was simply to deplete satellite cells in the experiments presented here. Immunostaining for Pax7 in muscles from *Sgcd*^*−/−*^; *Mapk3*^−/−^; *Mapk1*^*f/f-Pax7Cre-ER*^ mice showed a near complete absence of satellite cells (Figure 2C). The loss of satellite cells resulted in the complete lack of regeneration in muscle sections from *Sgcd*^*−/−*^; *Mapk3*^−/−^; *Mapk1*^*f/f-Pax7Cre-ER*^ mice as assessed by Myh3 positivity in myofibers and by counting myofibers with centrally located nuclei (Figure 2D-G). Skeletal muscles from *Sgcd*^*−/−*^; *Mapk3*^−/−^; *Mapk1*^*f/f-Pax7Cre-ER*^ mice were smaller than healthy and disease controls (Figure 2F and H). The total number of myofibers present in the extensor digitorum longus (EDL) muscle was counted at 8 weeks of age and *Sgcd*^*−/−*^; *Mapk3*^−/−^; *Mapk1*^*f/f-Pax7Cre-ER*^ mice had significantly fewer myofibers compared with controls (Figure 2I). We did not observe a significant change in the number of myofibers between wild type and *Sgcd*^*−/−*^; *Mapk3*^−/−^; *Mapk1*^*f/f*^ mice (Figure 2I). This reduction in muscle weights and total myofiber number in *Sgcd*^*−/−*^; *Mapk3*^−/−^; *Mapk1*^*f/f-Pax7Cre-ER*^ muscles is consistent with a loss of new myofiber formation due to deletion of satellite cells.

Unexpectedly, myofibers spared from degeneration in *Sgcd*^*−/−*^; *Mapk3*^−/−^; *Mapk1*^*f/f-Pax7Cre-ER*^ mice initiated a robust hypertrophic response in which fibers reached sizes never observed in wild type or disease controls (Figure 3A). At 8 weeks, interstitial fibrosis in the quadriceps of *Sgcd*^*−/−*^; *Mapk3*^−/−^; *Mapk1*^*f/f-Pax7Cre-ER*^ mice was reduced in comparison to *Sgcd*^*−/−*^; *Mapk3*^−/−^; *Mapk1*^*f/f*^ controls (Figure 3B-C). As readout for sarcolemma stability, we performed immunostaining for the immunoglobulin M (IgM) protein which is only present inside myofibers that have a compromised sarcolemma. A marked reduction in IgM positive myofibers was observed in muscle sections of *Sgcd*^*−/−*^; *Mapk3*^−/−^; *Mapk1*^*f/f-Pax7Cre-ER*^ quadriceps relative to those from *Sgcd*^*−/−*^; *Mapk3*^−/−^; *Mapk1*^*f/f*^ littermate controls (Figure 3D-E). Overall, our histological analyses reveal that, in the absence of satellite cell mediated regeneration, dystrophic myofibers show intrinsic protection from muscle damage and less pathologic features (Figure 2F). However, despite these dramatic histological improvements, these *Sgcd*^*−/−*^ mice in which satellite cells were deleted at 2 weeks of age (Figure 3F) eventually died likely due to an inability to generate new myofibers to replace lost ones in very early stages of MD (Figure 3G). *Sgcd*^*−/−*^; *Mapk3*^−/−^; *Mapk1*^*f/f-Pax7Cre-ER*^ mice had an average lifespan of 17.5 weeks while *Sgcd*^*−/−*^; *Mapk3*^−/−^; *Mapk1*^*f/f*^ lived beyond 40 weeks of age (Figure 3F-G). However, as will be discussed below, deleting satellite cells in early adulthood in dystrophic mice did not result in same lethality profile as mice in which satellite cells were deleted at 2 weeks of age.

**Figure 3.**
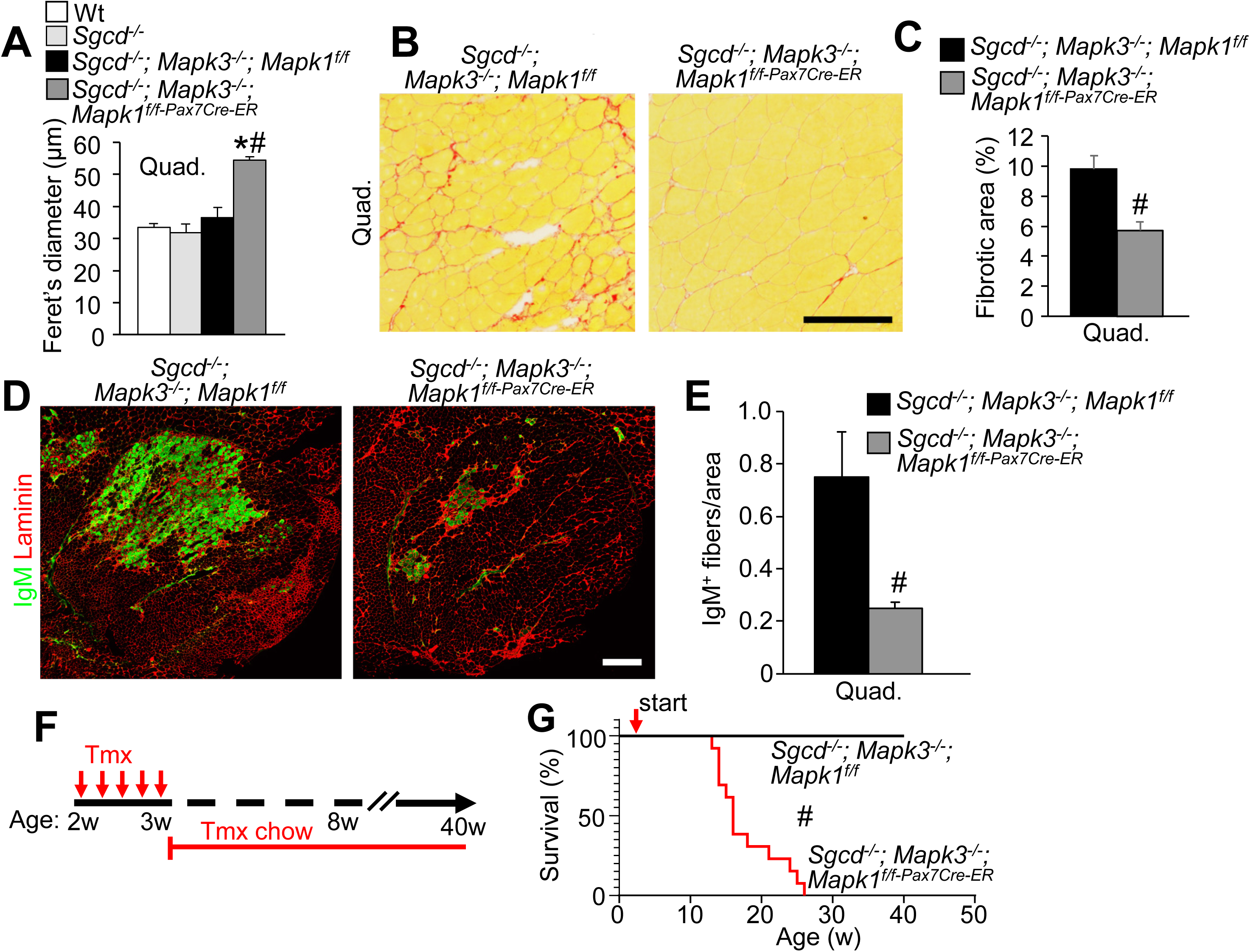
Loss of satellite cells reduces muscle damage in *Sgcd*^*−/−*^ mice. (A) Average minimal feret’s diameter values measured from muscle sections of the quad in mice with the indicated genotypes. A one-way ANOVA with Tukey’s multiple comparisons test was used to determine significance, *P < 0.05 versus wt, # P < 0.05 versus disease controls. (B) Representative Picrosirius Red-stained histological sections of the quad muscle from mice of the indicated genotypes. Scale bar = 100 μm. (C) Quantification of the fibrosis present in quad muscle sections from mice of the indicated genotypes. n = 3, *Sgcd*^*−/−*^; *Mapk3*^−/−^; *Mapk1^f/f^*; n = 4, *Sgcd*^*−/−*^; *Mapk3*^−/−^; *Mapk1*^*f/f-Pax7Cre-ER*^. Significance was determined using a Student’s t-test, #P < 0.05. (D) Representative histological sections of the quad muscle immunostained for immunoglobulin M (IgM) (green) and for laminin (Red) in mice of the indicated genotypes. Scale bar = 500 μm. (E) Quantification of IgM positive fibers in quad muscle sections from mice of the indicated genotypes. n = 5, *Sgcd*^*−/−*^; *Mapk3*^−/−^; *Mapk1^f/f^*; n = 6, *Sgcd*^*−/−*^; *Mapk3*^−/−^; *Mapk1*^*f/f-Pax7Cre-ER*^. Significance was determined using a Student’s t-test, #P < 0.05. (F) Schematic representation of the tmx treatment regimen. Two week-old mice received a daily tmx injection for 5 consecutive days and were subsequently placed on tmx chow after weaning. (G) Survival plot of mice with the indicated genotypes following tmx treatment starting at 2 weeks of age. #P < 0.05. Data represent mean ± SEM for A, C and E.

### Loss of satellite cells at peak disease reduces muscle damage in Sgcd^−/−^ mice

Next, we initiated tamoxifen treatment for deletion of *Mapk1/3* in satellite cells at 2 months of age, a time point when neonatal developmental hypertrophy of muscle is complete, but also a time point when ongoing MD pathology plateaus in *Sgcd*^*−/−*^ mice (Figure 4A). Mice were then harvested 2 months later, and quadriceps were assessed for satellite cell numbers (Figure 4A). Relative to wild type controls, *Sgcd*^*−/−*^ mice and control *Sgcd*^*−/−*^; *Mapk3*^−/−^; *Mapk1*^*f/f*^ mice had a relative expansion of satellite cells, which is typical of the dystrophic process in early adulthood in mice (Figure 4B). However, *Sgcd*^*−/−*^; *Mapk3*^−/−^; *Mapk1*^*f/f-Pax7Cre-ER*^ mice lacking *Mapk1/3* in their satellite cells showed a near complete loss of these cells (Figure 4B). This loss of satellite cells in *Sgcd*^*−/−*^; *Mapk3*^−/−^; *Mapk1*^*f/f-Pax7Cre-ER*^ mice was accompanied by a marked reduction in myofiber regeneration as very few myofibers were found to be Myh3 positive, a marker of embryonic myosin that is part of myofiber regeneration (Figure 4C-D). Relative to dystrophic controls, skeletal muscle weights from *Sgcd*^*−/−*^; *Mapk3*^−/−^; *Mapk1*^*f/f-Pax7Cre-ER*^ mice no longer showed pseudohypertrophy and were instead unchanged compared to wild type controls (Figure 4E). More importantly, histological analysis of quadriceps showed an improvement in gross pathologic features, such as fewer centrally nucleated myofibers and fewer small fibers, as well as an increase in larger fibers in the absence of *Mapk1/3* in satellite cells (Figure 4F and G). A significant decrease in total myofiber number in the EDL muscle was also observed in *Sgcd*^*−/−*^; *Mapk3*^−/−^; *Mapk1*^*f/f-Pax7Cre-ER*^ mice compared to *Sgcd*^*−/−*^; *Mapk3*^−/−^; *Mapk1*^*f/f*^ control mice, again suggesting that myofiber regeneration was extinguished (Figure 4H). Finally, there were also significantly fewer IgM positive myofibers in *Sgcd*^*−/−*^ muscle lacking satellite cells compared to muscle from *Sgcd*^*−/−*^ mice with satellite cells (Figure 4I). Thus, the loss of satellite cells at peak disease in *Sgcd*^*−/−*^ mice protected skeletal muscles from degeneration.

**Figure 4.**
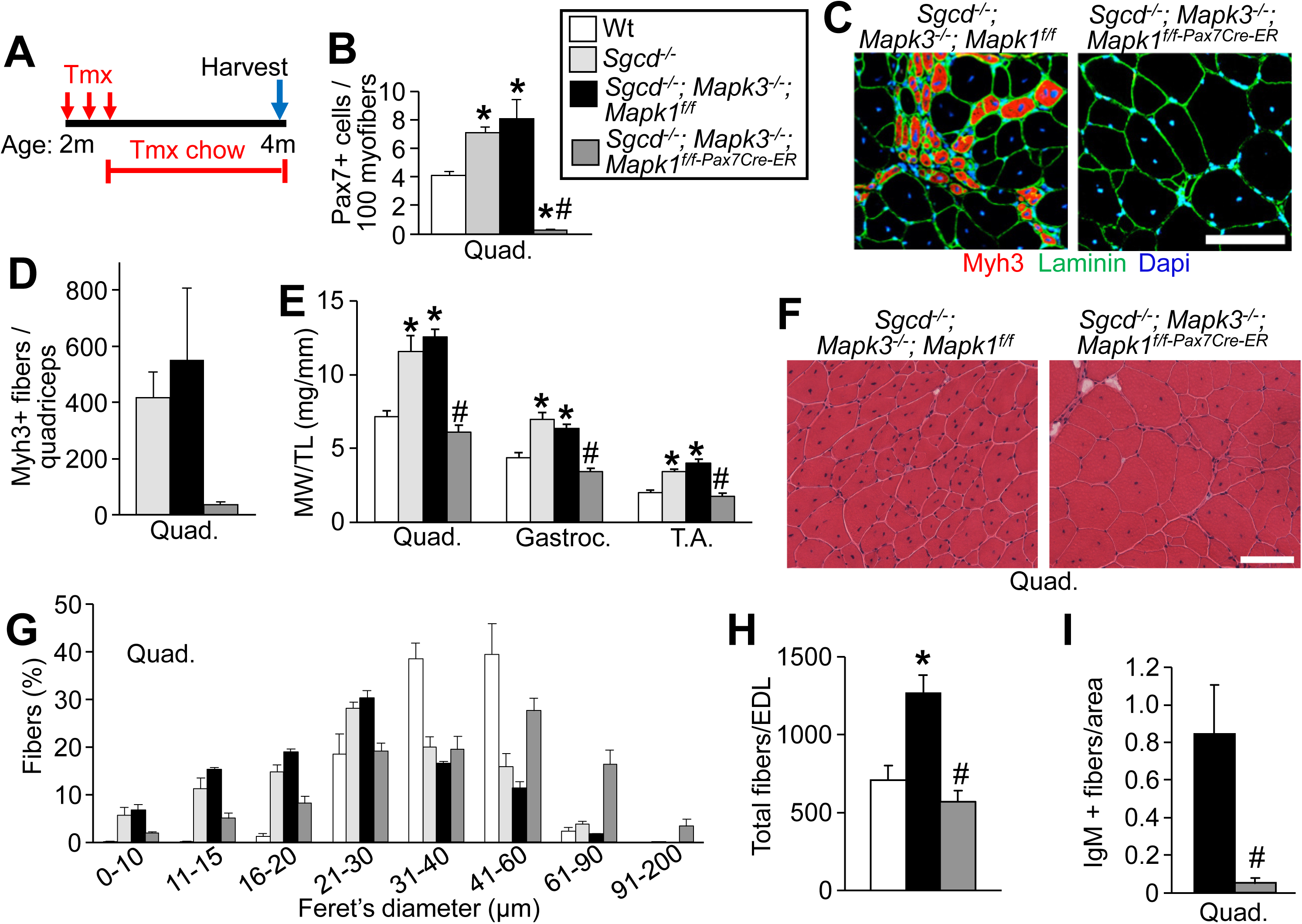
Satellite cell depletion at disease peak in *Sgcd*^*−/−*^ mice leads to improved histopathology. (A) Schematic representation of the tmx treatment regimen. Two-month-old mice received a daily tmx injection for 5 consecutive days and were subsequently placed on tmx chow after weaning. (B) Quantification of Pax7 positive satellite cells in muscle sections from the quad of mice with the indicated genotypes. n = 4 for all groups. A one-way ANOVA with Tukey’s multiple comparisons test was used to determine significance, *P < 0.05 versus wt, # P < 0.05 versus disease controls. (C) Representative quad muscle sections immunostained for Myh3 (red) and laminin (green) in 4-month-old mice of the indicated genotypes. Dapi stained nuclei are in blue. Scale bar = 100 μm. (D) Quantification of the number of Myh3 positive fibers in quad muscle sections of mice with the indicated genotypes. (E) M.W./T.L. ratios from mice of the indicated genotypes at 4 months of age. n = 8, wt; n = 4, *Sgcd*^*−/−*^; n = 6, *Sgcd*^*−/−*^; *Mapk3*^−/−^; *Mapk1^f/f^*; n = 6, *Sgcd*^*−/−*^; *Mapk3*^−/−^; *Mapk1*^*f/f-Pax7Cre-ER*^. A one-way ANOVA with Tukey’s multiple comparisons test was used to determine significance, *P < 0.05 versus wt, # P < 0.05 versus disease controls (F) Representative H&E stained sections of the quad muscle from 4-month-old mice of the indicated genotypes. Scale bar = 100 μm. (G) Minimal feret’s diameter distribution from the quad muscle of 4-month-old mice with the indicated genotypes. n = 4, wt; n = 4, *Sgcd*^*−/−*^; n = 3, *Sgcd*^*−/−*^; *Mapk3*^−/−^; *Mapk1^f/f^*; n = 3, *Sgcd*^*−/−*^; *Mapk3*^−/−^; *Mapk1*^*f/f-Pax7Cre-ER*^. (H) Quantification of the total myofibers present in the EDL muscle from 4-month-old mice of the indicated genotypes. n = 5 for all genotypes. A one-way ANOVA with Tukey’s multiple comparisons test was used to determine significance, *P < 0.05 versus wt, # P < 0.05 versus *Sgcd*^*−/−*^; *Mapk3*^−/−^; *Mapk1^f/f^*. (I) Quantification of IgM positive fibers in quad muscle section from mice of the indicated genotypes. n = 3, *Sgcd*^*−/−*^; *Mapk3*^−/−^; *Mapk1^f/f^*; n = 4, *Sgcd*^*−/−*^; *Mapk3*^−/−^; *Mapk1*^*f/f-Pax7Cre-ER*^. Data represent mean ± SEM for all graphs.

We next investigated whether the histological benefits observed with satellite cell deletion between 2-4 months of age were sustained over time. *Sgcd*^*−/−*^; *Mapk3*^−/−^; *Mapk1*^*f/f-Pax7Cre-ER*^ and littermate controls (*Sgcd*^*−/−*^; *Mapk3*^−/−^; *Mapk1^f/f^*) were treated with tamoxifen starting at 2 months of age and samples collected at 7 months of age (Supplemental Figure 3A). Similar to what was observed at the 4 month time point, there were fewer myofibers with centrally located nuclei at 7 months of age in muscle from *Sgcd*^*−/−*^ mice lacking satellite cells (Supplemental Figure 3B and C), and there were fewer IgM positive myofibers indicating greater myofiber membrane stability (Supplemental Figure 3D). Finally, we also measured the minimal feret’s diameter that again revealed a dramatic shift in myofiber size in *Sgcd*^*−/−*^; *Mapk3*^−/−^; *Mapk1*^*f/f-Pax7Cre-ER*^ muscle sections relative to controls (Supplemental Figure 3E). These data demonstrate that the intrinsic protective response in myofibers associated with the loss of satellite cells is sustained into mid-adulthood.

As a complementary approach, we also generated a satellite cell ablation mouse model in a dystrophic background using an inducible diphtheria toxin-based system. The *Pax7*^*CreER*^ mouse used to generate the *Mapk3*^−/−^; *Mapk1*^*f/f-Pax7Cre-ER*^ mice is very efficient so that when these mice are crossed with mice expressing the *Rosa26*-DTA (diphtheria toxin A) allele (Lepper et al., 2011) as a means of killing satellite cells, it is uniformly lethal. Because of this fact, we relied on another commonly used *Pax7*^*CreER*^ expressing mouse that did not cause lethality (Murphy et al., 2011). This *Pax7*^*CreER*^ knock-in mouse has an IRES-Cre cassette downstream of the endogenous stop codon, which we crossed with the Rosa26-DTA allele and treated 2-month-old mice with tamoxifen for 2 months (Supplemental Figure 4A and B). Relative to controls, *Sgcd*^*−/−*^; *R26-DTA*^*Pax7Cre-ER*^ had a significant reduction in satellite cells as assessed by immunostaining for Pax7 in histological sections (Supplemental Figure 4C). The decrease in satellite cells led to a parallel decrease in the number Myh3 positive regenerating myofibers (Supplemental Figure 4D and E). Muscles from 4-month-old *Sgcd*^*−/−*^; *R26-DTA*^*Pax7Cre-ER*^ mice were reduced in weight compared to *Sgcd*^*−/−*^ control but unchanged relative to wild type controls (Supplemental Figure 4F). An increase in the proportion of large caliber myofibers could be seen in *Sgcd*^*−/−*^; *R26-DTA*^*Pax7Cre-ER*^ quadriceps muscle sections as well as fewer myofibers with compromised sarcolemmal integrity as assessed by IgM immunostaining (Supplemental Figure 4G and H). The results obtained using *Sgcd*^*−/−*^; *R26-DTA*^*Pax7Cre-ER*^ mice are completely consistent with the results observed with *Mapk1/3* deletion, again showing that dystrophic myofibers lacking satellite cells hypertrophy and show relative protection from muscle damage and histopathology.

### Improved histopathology in mdx mice lacking satellite cells

To determine whether the loss of satellite cells could similarly reduce muscle pathology in a second dystrophic mouse model, we crossed *Mapk3*^−/−^; *Mapk1*^*f/f-Pax7Cre-ER*^ onto the *mdx*^*4Cv*^ background (Chapman et al., 1989), which more directly models DMD. We treated *mdx*; *Mapk3*^−/−^; *Mapk1*^*f/f-Pax7Cre-ER*^ and *mdx* control mice with tamoxifen starting at 2 weeks of age prior to disease onset (Figure 5A). *mdx* mice were then harvested 6 weeks later at 8 weeks of age and we again observed a dramatic depletion of satellite cells in *mdx*; *Mapk3*^−/−^; *Mapk1*^*f/f-Pax7Cre-ER*^ muscle sections compared to *mdx* control (Figure 5B). These *mdx* mice lacking satellite cells also showed an absence of Myh3 positive myofibers and essentially no myofibers with centrally located nuclei (Figure 5C-E). Histopathology was again noticeably improved and myofiber diameters were dramatically enhanced in *mdx*; *Mapk3*^*−/−*^; *Mapk1*^*f/f-Pax7Cre-ER*^ relative to wild type and *mdx* controls (Figure 5E-F).

**Figure 5.**
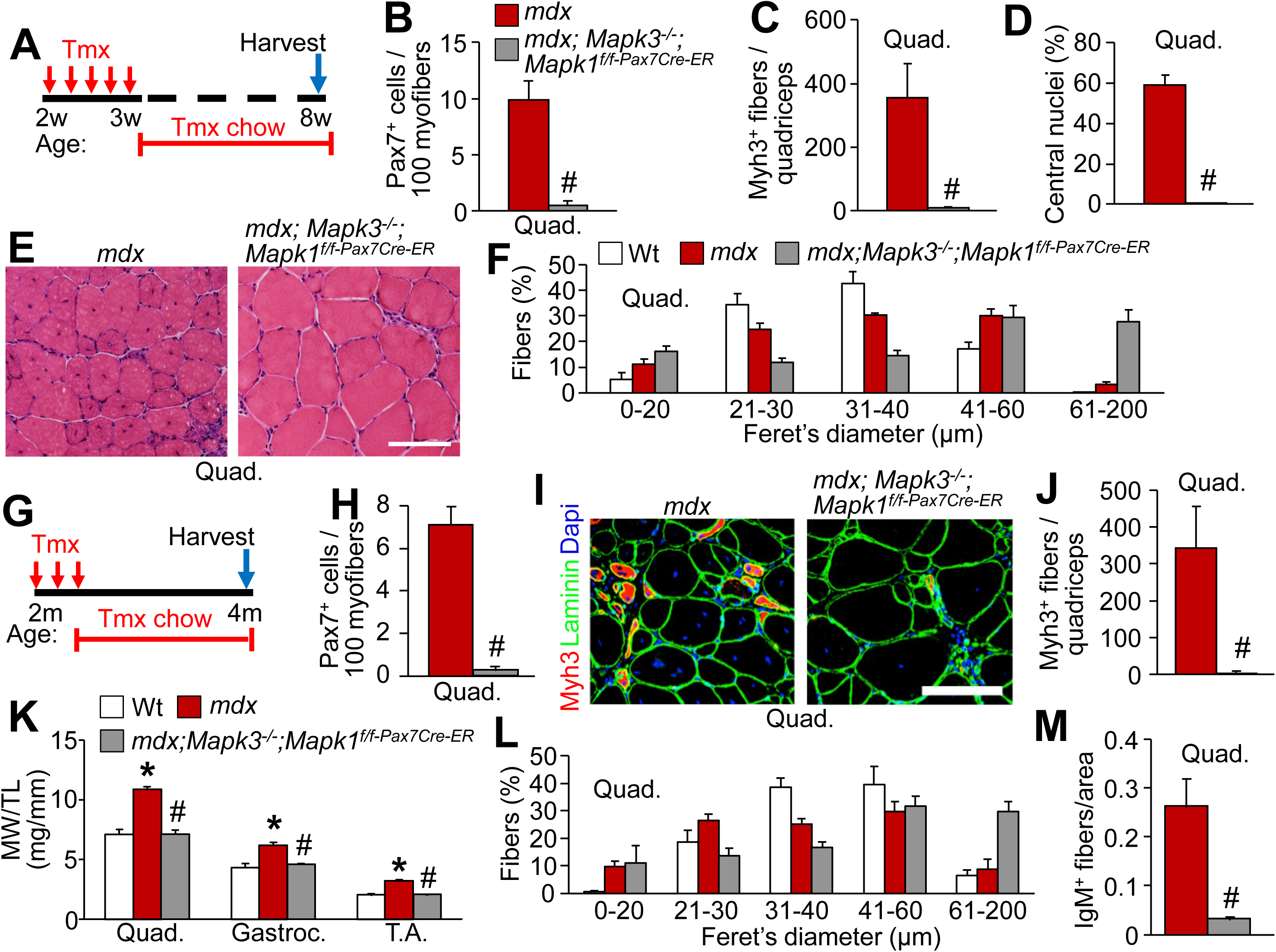
Improved histopathology in *mdx* mice lacking satellite cells. (A) Schematic representation of the tmx treatment in *mdx* mice. Two week-old mice received a daily tmx injection for 5 consecutive days and were subsequently placed on tmx chow after weaning. (B) Quantification of satellite cell numbers from quad muscle sections from 2-month-old mice of the indicated genotypes. n = 3 for both groups. Significance was determined using a Student’s t-test, #P < 0.05. (C) Quantification of the number of Myh3 positive fibers in quad muscle sections of mice with the indicated genotypes. n = 3, *mdx*; n = 4, *mdx*; *Mapk3*^−/−^; *Mapk1*^*f/f-Pax7Cre-ER*^. Significance was determined using a Student’s t-test, #P < 0.05. (D) Quantification of the number of myofibers with centrally located nuclei in quad muscle sections from mice of the indicated genotypes. n = 3, *mdx* n = 4, *mdx*; *Mapk3*^−/−^; *Mapk1*^*f/f-Pax7Cre-ER*^. Significance was determined using a Student’s t-test, # P < 0.05. (E) Representative H&E stained histological sections of the quad muscle from mice of the indicated genotypes. Scale bar = 100 μm. (F) Minimal feret’s diameter distribution from the quad muscle of 2-month-old mice with the indicated genotypes. n = 4 for all groups. The same 2-month wt weight data are also shown in Figure 2. (G) Schematic representation of the tmx treatment regimen. Two-month-old mice received a daily tmx injection for 5 consecutive days and were subsequently placed on tmx chow after weaning. (H) Quantification of satellite cell numbers from quad muscle sections from 4-month-old mice of the indicated genotypes. n = 4, *mdx* n = 3, *mdx*; *Mapk3*^−/−^; *Mapk1*^*f/f-Pax7Cre-ER*^. Significance was determined using a t-test, #P < 0.05. (I) Representative quad muscle sections immunostained for Myh3 (red) and laminin (green) in 4-month-old mice of the indicated genotypes. Dapi stained nuclei are in blue. Scale bar = 100 μm. (J) Quantification of the number of Myh3 positive fibers in quad muscle sections of mice with the indicated genotypes. n = 3 for both groups. Significance was determined using a Student’s t-test, #P < 0.05. (K) M.W./T.L. ratios from mice of the indicated genotypes at 4 months of age. n = 5 for all groups. A one-way ANOVA with Tukey’s multiple comparisons test was used to determine significance, *P < 0.05 versus wt, # P < 0.05 versus *mdx*. The 4-month wt weight data are also shown in Figure 4. (L) Minimal feret’s diameter distribution from the quad muscle of 4-month-old mice with the indicated genotypes. n = 4, wt; n = 3, *mdx* n = 3, *mdx*; *Mapk3*^−/−^; *Mapk1*^*f/f-Pax7Cre-ER*^. The 4-month wt feret’s diameter data are also shown in Figure 4. (M) Quantification of IgM positive fibers in quad muscle sections from mice of the indicated genotypes. n = 5 for both groups. Significance was determined using a Student’s t-test, #P < 0.05. Data represent mean ± SEM for all graphs.

We also assessed whether depleting satellite cells beginning at 2 months of age, after neonatal development in *mdx* mice would similarly provide protection (Figure 5G). Satellite cells were again completely eliminated and Myh3 positive de novo myofibers were absent in 4-month-old *mdx*; *Mapk3*^−/−^; *Mapk1*^*f/f-Pax7Cre-ER*^ mice relative to controls (Figure 5H-J). Muscle weights of *mdx*; *Mapk3*^−/−^; *Mapk1*^*f/f-Pax7Cre-ER*^ mice were similar to wild type controls but significantly smaller than the pseudohypertrophy observed in *mdx* controls (Figure 5K). Compared to wild type and *mdx* controls, a greater proportion of large caliber myofibers was observed in 4-month-old *mdx*; *Mapk3*^−/−^; *Mapk1*^*f/f-Pax7Cre-ER*^ muscle sections of the quadriceps (Figure 5L). Finally, as we described in *Sgcd*^*−/−*^ mice lacking satellite cells, a significant reduction in IgM positive myofibers was observed in histological sections of *mdx* mice without satellite cells compared to those with satellite cells (Figure 5M). Hence, ablation of satellite cells in a mouse model of DMD, the most common form of MD, yielded a similar protective beneficial response as observed in *Sgcd*^*−/−*^ mice.

### Identification of an adaptive window in dystrophic mice

To better understand the protective window whereby ablating satellite cells could provide benefit to ongoing disease pathogenesis in *Sgcd*^*−/−*^ mice we treated 6-month-old mice. Here we again observed a robust deletion of satellite cells from 6-8 months of age in *Sgcd*^*−/−*^; *Mapk3*^−/−^; *Mapk1*^*f/f-Pax7Cre-ER*^ mice, which was associated with a virtual absence of Myh3 positive myofibers (Supplemental Figure 5A-C). The absence of satellite cells in *Sgcd*^*−/−*^ mice at 8 months of age also led to a reduction in the number of myofibers and muscle weights relative to *Sgcd*^*−/−*^ mice with satellite cells (Supplemental Figure 5D and E). However, at this later time point the absence of satellite cells did not skew the myofiber size distribution towards larger values in the quadriceps of *Sgcd*^*−/−*^ mice, and tissue histopathology appeared rather similar (Supplemental Figure 5F and G). The lack of myofiber hypertrophy in *Sgcd*^*−/−*^; *Mapk3*^−/−^; *Mapk1*^*f/f-Pax7Cre-ER*^ mice and the lack of noticeable improvement in tissue histopathology may simply reflect the disease process from the first 6 months of life in these *Sgcd*^*−/−*^ mice that persists, or that the effect of satellite cell ablation is less effective in more mature dystrophic muscle. Thus, it is likely that we have delineated a time window in which myofibers most favorably adapt to chronic dystrophic disease due to satellite cell deletion.

### Improved running capacity in Sgcd^−/−^ lacking satellite cells

To test whether the improved histopathology associated with satellite cell depletion in *Sgcd*^*−/−*^ mice enhanced muscle function, we challenged mice to run on a treadmill as an indirect readout of muscle performance. Mice were subjected to forced downhill running with a ramping speed protocol (Goonasekera et al., 2011). We depleted satellite cells in *Sgcd*^*−/−*^ mice prior to disease onset by treating the mice with tamoxifen starting at 2 weeks of age (Figure 6A). On average, *Sgcd*^*−/−*^ mice ran half the distance and had a greater number of shock pad visits compared to wild type controls (Figure 6B-C). However, consistent with the mitigated histopathology, *Sgcd*^*−/−*^ mice lacking satellite cells performed significantly better that those with satellite cells, albeit not to wild type standards (Figure 6B-C). Depleting satellite cells after the neonatal development period by giving tamoxifen between 2-4 months of age also significantly rescued treadmill running performance in *Sgcd*^*−/−*^; *Mapk3*^−/−^; *Mapk1*^*f/f-Pax7Cre-ER*^ mice compared to to disease controls (Figure 6D-F). In conclusion, the lack of satellite cells in skeletal muscles of *Sgcd*^*−/−*^ mice in early adulthood is associated with fewer damaged myofibers and improved running performance.

**Figure 6.**
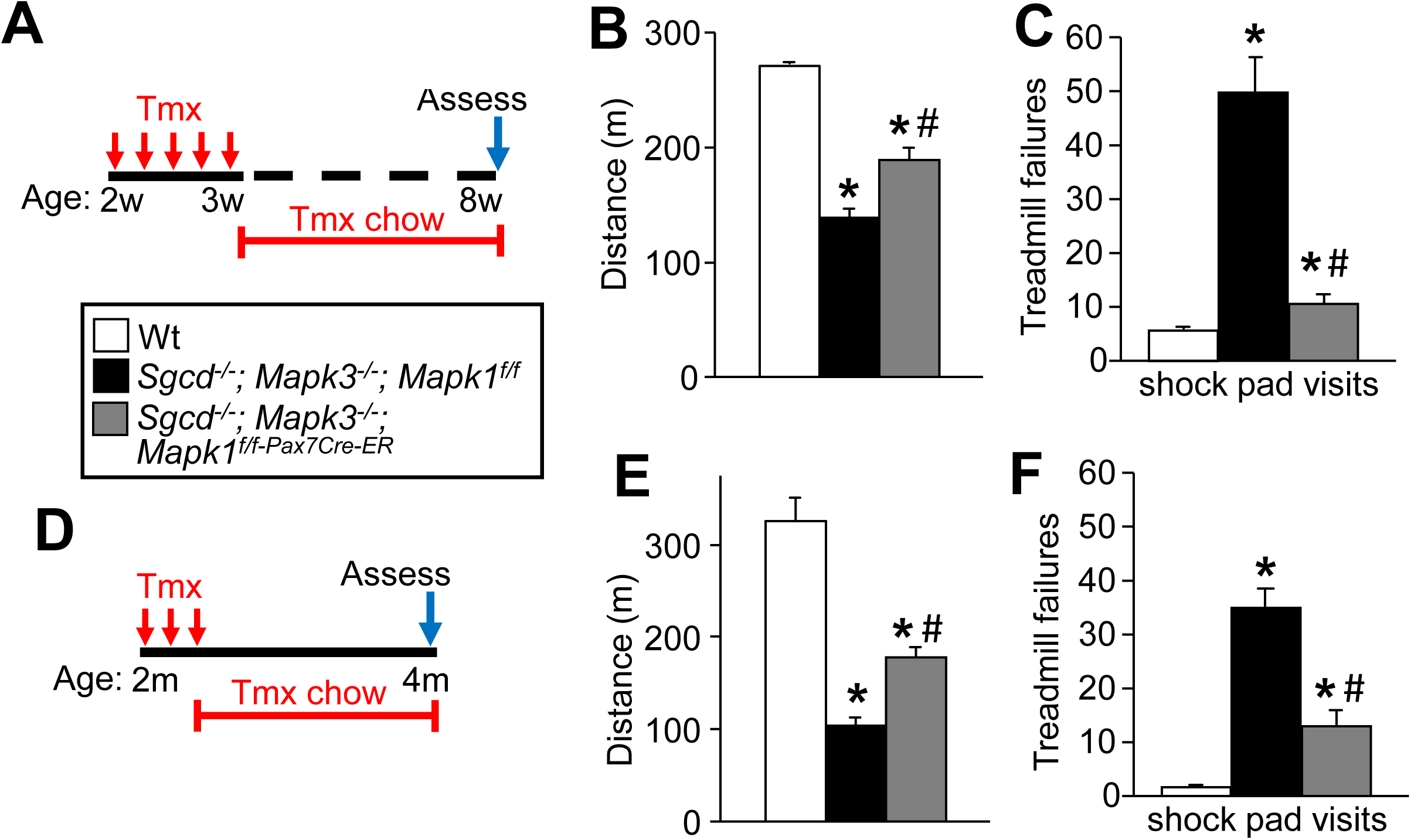
Improved running capacity in *Sgcd*^*−/−*^ mice lacking satellite cells. (A) Schematic representation of the tmx treatment regimen. Two-week-old mice received a daily tmx injection for 5 consecutive days and were subsequently placed on tmx chow after weaning. (B) Average forced running distance on a treadmill for 2-month-old mice of the indicated genotypes. n = 4, wt; n = 5, *Sgcd*^*−/−*^; *Mapk3*^−/−^; *Mapk1^f/f^*; n = 3, *Sgcd*^*−/−*^; *Mapk3*^−/−^; *Mapk1*^*f/f-Pax7Cre-ER*^. A one-way ANOVA with Tukey’s multiple comparisons test was used to determine significance, *P < 0.05 versus wt, # P < 0.05 versus *Sgcd*^*−/−*^; *Mapk3*^−/−^; *Mapk1^f/f^*. (C) Quantification of the number of shock pad visits calculated over 20 min from the start of the protocol. n = 4, wt; n = 5, *Sgcd*^*−/−*^; *Mapk3*^−/−^; *Mapk1^f/f^*; n = 3, *Sgcd*^*−/−*^; *Mapk3*^−/−^; *Mapk1*^*f/f-Pax7Cre-ER*^. A one-way ANOVA with Tukey’s multiple comparisons test was used to determine significance, *P < 0.05 versus wt, # P < 0.05 versus *Sgcd*^*−/−*^; *Mapk3*^−/−^; *Mapk1^f/f^*. (D) Schematic representation of the tmx treatment regimen. Two-month-old mice received a daily tmx injection for 5 consecutive days and were subsequently placed on tmx chow after weaning. (E) Average forced running distance on a treadmill for 4-month-old mice of the indicated genotypes. n = 4, wt; n = 3, *Sgcd*^*−/−*^; *Mapk3*^−/−^; *Mapk1^f/f^*; n = 5, *Sgcd*^*−/−*^; *Mapk3*^−/−^; *Mapk1*^*f/f-Pax7Cre-ER*^. A one-way ANOVA with Tukey’s multiple comparisons test was used to determine significance, *P < 0.05 versus wt, # P < 0.05 versus *Sgcd*^*−/−*^; *Mapk3*^−/−^; *Mapk1^f/f^*. (F) Quantification of the number of shock pad visits calculated 10 min from the start of the protocol. n = 4, wt; n = 3, *Sgcd*^*−/−*^; *Mapk3*^−/−^; *Mapk1^f/f^*; n = 5, *Sgcd*^*−/−*^; *Mapk3*^−/−^; *Mapk1*^*f/f-Pax7Cre-ER*^. A one-way ANOVA with Tukey’s multiple comparisons test was used to determine significance, *P < 0.05 versus wt, # P < 0.05 versus *Sgcd*^*−/−*^; *Mapk3*^−/−^; *Mapk1^f/f^*. Data represent mean ± SEM for all graphs.

## Discussion

Muscle degeneration, the pathological hallmark of MDs, is followed by satellite cell-mediated regeneration of de novo myofibers, although these newly myofibers are still susceptible to subsequent necrosis because they harbor the disease-causing mutation. While the skeletal muscle field assumes that satellite cell-mediated regeneration is a strictly beneficial compensatory response in MD, this has never been directly examined in vivo. However, 2 previous studies provided an indirect assessment of this concept in *mdx* mice. Granata et al. (1998) along with Pagel and Partridge (1999) used irradiation of one limb to inactivate satellite cells and compared disease pathogenesis to the non-irradiated control leg. The results showed that muscle histopathology was noticeably improved when satellite cells were inactivated, suggesting for the first time that perhaps satellite cells were not strictly beneficial during MD (Granata et al., 1998). These observations from decades ago were never extended, although here we employed multiple independent lines of genetic investigation that each suggest satellite cell activation during young adulthood appears to do more harm than good.

Here we show that *Mapk3*^−/−^; *Mapk1*^*f/f-Pax7Cre-ER*^ mice serve as a novel and highly efficient cell ablation model to examine the in vivo relevance of satellite cells in MD. We showed that dystrophic mice lacking satellite cells were not only able to survive the initial phase of muscle degeneration; they initiated an intrinsic protective response against muscle damage in myofibers. Nevertheless, despite the improved histopathology in dystrophic mice lacking satellite cells, eliminating these cells during early neonatal development (beginning at 2 weeks of age) did reduce their lifespan in an MD context. However, this is a complex situation because the mice have not yet attained a critical mass of muscle to sustain adult function (White et al., 2010), and these mice are undergoing their first major round of muscle degeneration that begins at 3 weeks of age (DiMario et al., 1991; Duddy et al., 2015). Deletion of satellite cells beginning at 8 weeks of age, when neonatal developmental maturation of muscle is mostly complete, and both *mdx* and *Sgcd*^*−/−*^ mice have already progressed through the most severe degeneration cycles, now appears to provide a much more sustained benefit to muscle histopathology and function compared to dystrophic mice replete in satellite cell *Mapk1/3*.

The underlying molecular mechanism whereby satellite cells ablation protects existing myofibers in dystrophic mice is uncertain. We observed a complete lack of new fiber formation in both *mdx* and *Sgcd*^*−/−*^ mice upon *Mapk1/3* deletion in satellite cells, which then produced a compensatory response that made existing myofibers dramatically larger. However, small caliber type I myofibers are believed to be less vulnerable to degeneration (Deconinck and Dan, 2007; Karpati et al., 1988), while the largest fibers appear to be more susceptible to contraction-induced damage in DMD (Lieber and Friden, 1988; Webster et al., 1988). Consistent with these observeations, increasing muscle mass in MD has been explored as a therapeutic approach. Specifically, inhibition of myostatin, a negative regulator of muscle mass, led to strength increases in *mdx* mice and better performance (Bogdanovich et al., 2002; Jin et al., 2019; St Andre et al., 2017). However, ultimately inhibition of myostatin in *mdx* mice failed to protect muscles from eccentric contraction-induced damage, thus we believe it is unlikely that the larger fibers observed here mechanistically explain the profound reduction in myofiber sarcolemmal rupture events and attenuated disease features.

However, our working hypothesis is that satellite cell activity itself could be additionally destabilizing to myofiber membranes of dystrophic mice through the fusion process. While satellite cell fusion is meant to revitalize damaged myofibers following acute injury, we really lack an understanding of how this process impacts a chronic disease like MD. Indeed, it is unlikely that satellite cells evolved to deal with chronic wasting diseases like MD where skeletal muscles show inherent sarcolemmal fragility. Satellite cells likely evolved as a means of continuing muscle growth during neonatal development and to replace muscle in an adult following an acute injury. In these 2 states, myofibers that are being invaded by new myoblasts from the satellite cells are not defective and can likely accommodate this event without detriment. However, the sarcolemma of dystrophic myofibers is destabilized and perhaps the very process of myoblast fusion exacerbates ongoing disease. During MD, satellite cells appear to fuse at focal sites of injury as opposed to a more diffuse fusion response (Blaveri et al., 1999). In turn, the volume of fusion along the myofiber may contribute to an unbalanced destabilization of the sarcolemma in MD. Our observations show that without satellite cells existing myofibers essentially lack membrane rupture events.

Our data are most consistent with a working model whereby slowing the regenerative response in MD, either by reducing the satellite cell pool or by delaying satellite function, may offer an advantage to existing myofibers during late muscle development.

## Materials and Methods

### Animal Models

Animal experiments performed in the study were approved by the Institutional Animal Care and Use Committee of the Cincinnati Children’s Hospital Medical Center. Mice did not undergo randomization because they were genetically identical and many groups were from the same litters, as well as matched for age and sex ratio. Both male and female mice were used in an equal ratio, and no sex-specific differences were observed.

Mice containing a genetic insertion of a tamoxifen-inducible Cre recombinase–estrogen receptor fusion protein, CreER^T2^ cDNA into the Pax7 *locus* (Lepper et al., 2009) or transgenic Ska-MCM mice (Boyer et al., 2019) were crossbred with *Mapk3*^*−/−*^ (Erk1) (Pages et al., 1999) and *Mapk1*^*f/f*^ (Erk2) mice (Samuels et al., 2008). *Mapk3*^−/−^; *Mapk1*^*f/f-Pax7Cre-ER*^ were crossed with *Rosa26* loxP site– dependent reporter mice (*R26eGFP*) to visualize satellite cells as well as onto the *Sgcd*^*−/−*^ (Hack et al., 2000) or the *mdx*^*4Cv*^ (Chapman et al., 1989) backgrounds to study the effects of satellite cell depletion in dystrophic mouse models. Other mouse lines used in the study were: *Rosa26-DTA* (R26-DTA, Jax stock No:006331) (Ivanova et al., 2005) and *Pax7*^*CreERT2*^ (Murphy et al., 2011) mice.

### Animal Procedures

Tamoxifen was administered to all experimental and control mice via intraperitoneal injections at (75 mg/kg) using pharmaceutical-grade tamoxifen (Sigma) dissolved in corn oil (Sigma). Subsequently, in experiments requiring longer term tamoxifen regimens, the mice were fed a diet containing 400 mg/kg tamoxifen citrate (Envigo, TD.55125) for the indicated time. Muscle injury was induced by injecting the TA muscle with 50 μl of cardiotoxin (10 μmol/L).

### Western Blotting

Muscles were homogenized and lysates prepared as previously described (Wissing et al., 2014). Proteins were resolved by SDS-PAGE, transferred to Immobilon-FL membranes (Millipore) and incubated overnight with antibodies against, phosphorylated Erk1/2 (1:1000, Cell Signaling Technology), Erk1/2 (1:2000, Cell Signaling Technology) and GAPDH (1:500 000, Fitzgerald), Membranes were then incubated with IRDye secondary antibodies (1:6000, LI-COR Biosciences) and visualized using an Odyssey CLx imaging system (LI-COR Biosciences).

### Immunofluorescence

Histological sections (8 μm) were collected from frozen skeletal muscles using a cryostat and fixed in 4% paraformaldehyde (PFA) for 10 min. Sections were subsequently washed in phosphate buffer saline (PBS) and heated in citrate buffer (pH 6.0) for 20 min and maintained in blocking buffer for 30 min (10% goat serum and 0.4% triton X diluted in PBS). Slides were stained overnight at 4°C with antibodies diluted in staining solution (1% bovine serum albumin, 0.04% triton X diluted in PBS). Satellite cells were visualized by staining for Pax7 (1:10, Developmental Studies Hybridoma Bank (DSHB)) and regenerating myofibers were identified by staining for Myh3 (F1.652, DSHB). Myofibers present in a muscle section were stained with an anti-myosin antibody (MF 20, DSHB) while anti-laminin antibody (Sigma) was used to visualize the outline of all myofibers present in a given muscle section. Nuclei were visualized using Dapi (Invitrogen). IgM primary antibody conjugated to FITC (1:300, Sigma) was used to highlight myofibers with compromised membrane integrity. TA muscles analyzed for GFP reporter activity were fixed in 4% PFA for 4 hrs at 4°C and then washed in PBS and equilibrated overnight in 30% sucrose diluted in PBS. Sections were stained for GFP (Novus).

Primary antibodies were visualized using Alexa-568 and 488 goat anti-rabbit IgG (Invitrogen), Alexa-488 goat anti-mouse IgG2b (Invitrogen) and Alexa-568 anti-mouse IgG1 (Invitrogen) secondary antibodies diluted 1:500 in staining solution. Immunofluorescence images were captured using a Nikon Eclipse Ti microscope or a Nikon A1R confocal microscope using NIS Elements AR 4.13 software. Data relating to satellite cell counts were generated by counting Pax7 positive cells relative to each myofiber present in two 10x images. Regenerating myofibers were assessed by counting Myh3 positive myofibers from an entire muscle section at mid-belly for a given animal. IgM positive myofibers were counted from entire muscle sections taken at the mid-belly of the muscle and normalized to muscle section area. All analyses were performed in a blinded fashion whereby the experimenter was only made aware of the genotypes following proof of quantification.

### Pathological indices

Histological cross-sections (8 μm) were collected from skeletal muscles using a cryostat and stained for H&E or Picrosirius Red to assess myofibers with centrally located nuclei and fibrosis, respectively. The number of myofibers with centrally located nuclei was quantified from two 10x micrographs taken from histological sections of the quadriceps at mid-belly. Fibrosis was quantified from two 10x pictures taken from histological sections of the quadriceps. Quantification of fibrotic area was calculated using the ImageJ analysis software. The minimal feret’s diameter was determined from laminin stained muscle sections using ImageJ. All analyses were performed in a blinded fashion whereby the experimenter was only made aware of the genotypes following proof of quantification.

### Fluorescence-activated cell sorting (FACS) and qRT-PCR

Skeletal muscles from tamoxifen treated mice were isolated, minced and digested in DMEM containing collagenase D (0.75U/ml, Roche) and dispase II (1.0U/ml, Roche). Once digested, the cells were passed through a cell strainer (70 μm) and pelleted down. Cells were then treated with red blood lysing buffer (Sigma) before being collected by fluorescence-activated cell sorting (FACS) for the endogenous eGFP fluorescence expressed by the recombined *R26eGFP* reporter allele. RNA was extracted from FACS-sorted isolated satellite cells using a Qiashredder homogenization instrument (Qiagen) and the RNAeasy kit according to the manufacturer’s instructions (Qiagen). Total RNA was reverse transcribed using random oligo-dT primers with a Verso cDNA synthesis kit (Thermo Fisher Scientific) according to manufacturer’s instructions. Quantitative real-time PCR was performed using Sso Advanced SYBR Green (Bio-Rad). ΔΔCT was used to quantify the fold change of the target genes. The following primer set was used to identify the *Mapk1* transcript: 5’ - TTACTCTACTTCTCCCCACTCC and 5’ - CTGCCTCTGACTTCTGAATG. GAPDH was used as an internal control 5′-TGACCACAGTCCATGCCATC and 5′-GACGGACACATTGGGGGTAG.

### Treadmill running

Mice were subjected to forced down-hill treadmill running using a ramping speed protocol as previously described (Goonasekera et al., 2011). Mice were run until exhaustion or until the entire protocol was completed.

### Statistics

A one-way analysis of variance was used to determine if there was a significant difference in experiments with more than 2 groups; a Tukey’s post hoc test was performed to compare individual groups (Prism Software). Significant differences between 2 groups were determined using an unpaired Student’s *t* test (Stats Plus Software). Significance was set at P<0.05 for all experiments. All results are presented as mean and the error bar represents the standard error of the mean.

## Acknowledgements

J.D.M. was funded by grants from the National Heart Lung and Blood Institute of the National Institutes of Health, and by the Howard Hughes Medical Institute. J.G.B. was funded by a grant from the Muscular Dystrophy Association, and R.J.V. was funded by Career Development Award From the American Heart Association

