## Supplemental for "Satellite cell depletion in early adulthood attenuates muscular dystrophy pathogenesis"

**SUPPLEMENTAL FIGURES 1-5**

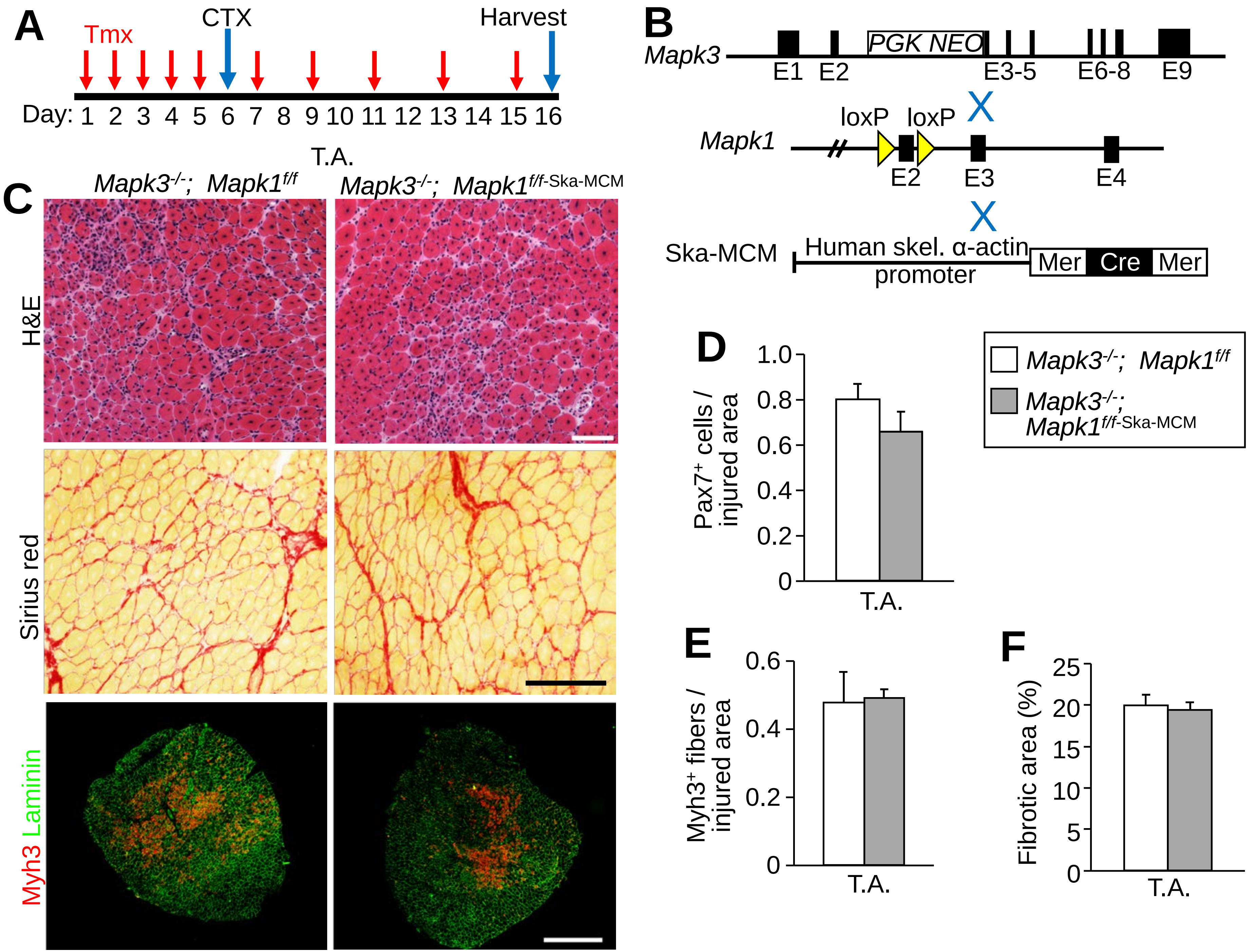

A
Tmx
CTX
Harvest
Day: 1 2 3 4 5 6 7 8 9 10 11 12 13 14 15 16
B
Mapk3
PGK NEO
E1 E2 E3-5 E6-8 E9
Mapk1
loxP loxP
E2 E3 E4
Ska-MCM
Human skel. α-actin promoter
Mer Cre Mer
T.A.
C
Mapk3-/-; Mapk1f/f
Mapk3-/-; Mapk1f/f-Ska-MCM
H&E
Sirius red
Myh3 Laminin
D
Pax7+ cells / injured area
1.0 0.8 0.6 0.4 0.2 0
T.A.
Mapk3-/-; Mapk1f/f
Mapk3-/-; Mapk1f/f-Ska-MCM
E
Myh3+ fibers / injured area
0.6 0.4 0.2 0
T.A.
F
Fibrotic area (%)
25 20 15 10 5 0
T.A.

**Supplemental Figure 1**. Loss of *Mapk3* and *Mapk1* during differentiation does not impact muscle regeneration. (A) Schematic showing the timing of tamoxifen (tmx) and cardiotoxin (CTX) treatments in 2-month-old mice. (B) Schematic representation of breeding of *Mapk3*$^{-/-}$ and *Mapk1*-loxP-targeted mice with Ska-MCM transgenic mice. (C) Representative H&E (top) and Picrosirius Red (middle)-stained sections of the tibialis anterior (T.A.) muscle 10 days post CTX injury. Scale bars = 100 µm. Immunohistochemistry (bottom panels) for Myh3 (red) and laminin (green) in T.A. muscle sections from mice of the indicated genotypes. Scale bar = 500 µm. (D) Quantification of Pax7 positive cells 10 days following CTX injury in T.A. muscle sections from mice of the indicated genotypes. n = 6, *Mapk3*$^{-/-}$; *Mapk1*$^{f/f}$; n = 5, *Mapk3*$^{-/-}$; *Mapk1*$^{f/f\text{-}Ska\text{-}MCM}$. (E) Quantification of Myh3 positive myofibers 10 days following CTX injury in T.A. muscle sections from mice of the indicated genotypes. n = 6, *Mapk3*$^{-/-}$; *Mapk1*$^{f/f}$; n = 5, *Mapk3*$^{-/-}$; *Mapk1*$^{f/f\text{-}Ska\text{-}MCM}$. (F) Quantification of the fibrosis present in T.A. muscle sections from mice of the indicated genotypes 10 days following CTX injury. n = 6, *Mapk3*$^{-/-}$; *Mapk1*$^{f/f}$; n = 5, *Mapk3*$^{-/-}$; *Mapk1*$^{f/f\text{-}Ska\text{-}MCM}$. Data represent mean ± SEM for all graphs.

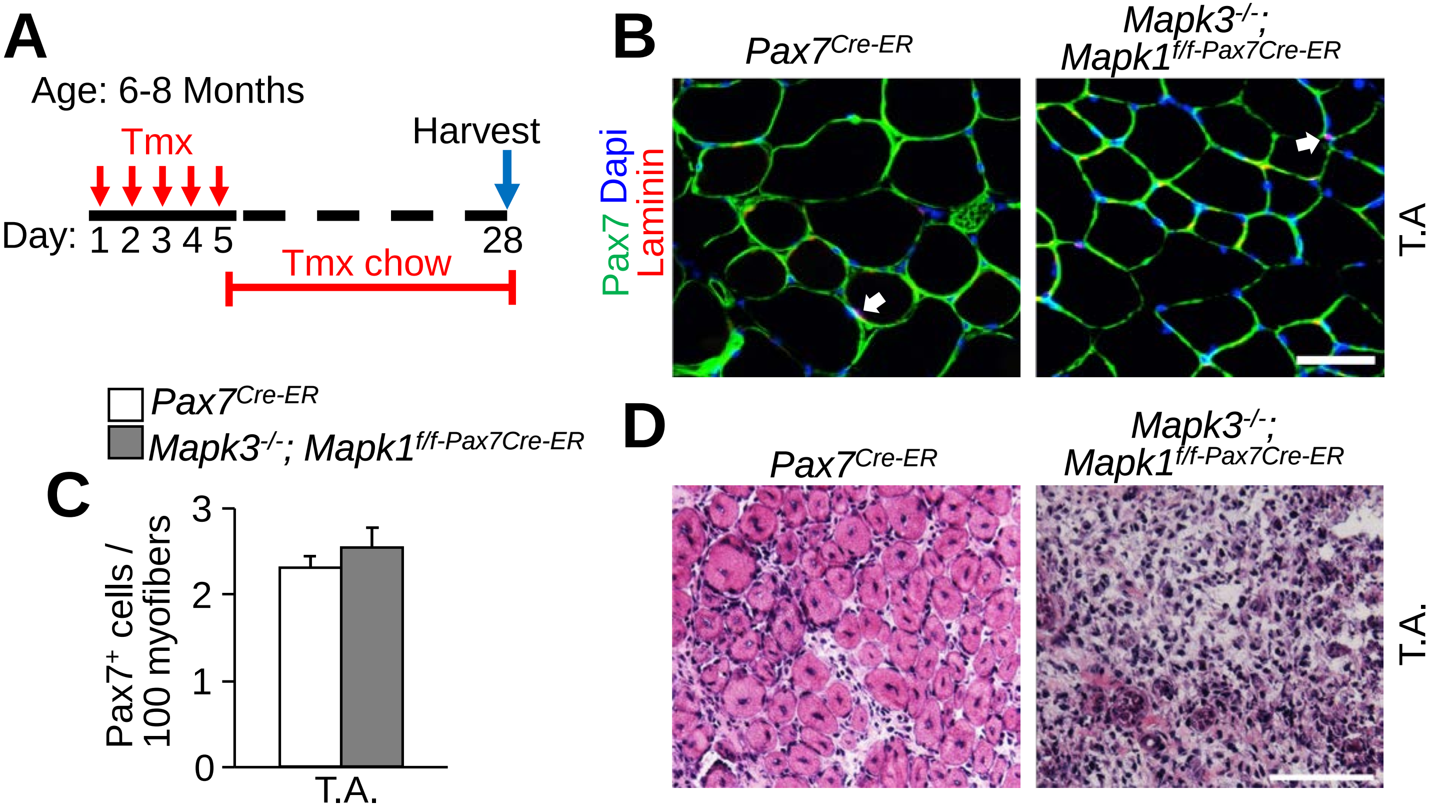


**Supplemental Figure 2**. Normal satellite cells numbers in adult mice lacking *Mapk3* and *Mapk1*. (A) Schematic representation of the tamoxifen (tmx) treatment regimen. Adult mice (6-8-month-old) received a daily tmx injection for 5 consecutive days and were subsequently placed on tmx chow after for 3 weeks. (B) Representative immunostained sections of the tibialis anterior (T.A.) muscle for Pax7 (arrows) and laminin (green). Dapi stained nuclei are in blue. Scale bar = 50 μm. (C) Quantification of Pax7 positive cells in T.A. muscle sections from mice of the indicated genotypes. n = 6 for both groups. Data represent mean ± SEM. (D) Representative H&E stained sections of the T.A. muscle 7 days post CTX injury. Scale bar = 100 μm.

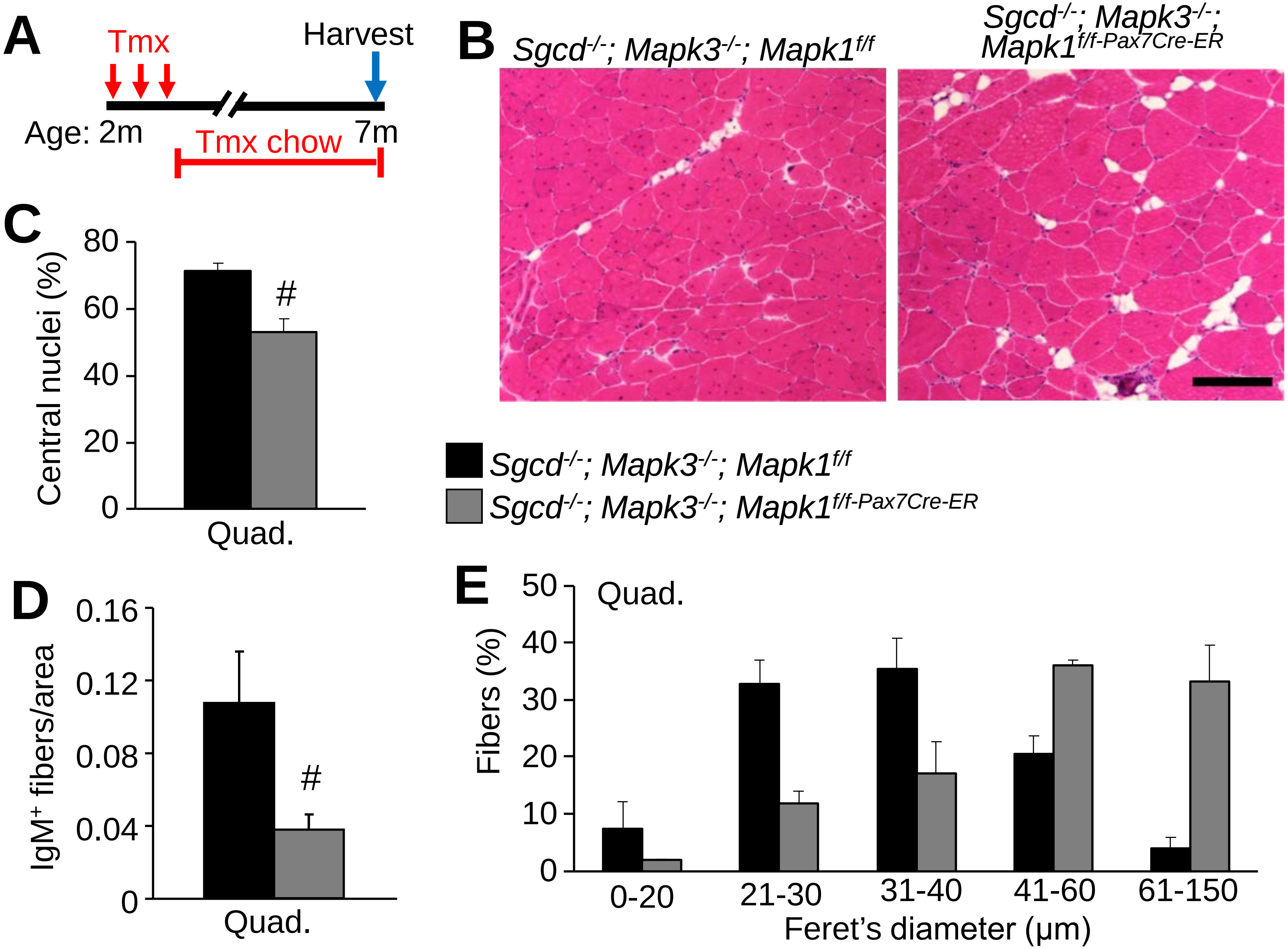
A
Tmx
Harvest
Age: 2m
7m
Tmx chow
B
Sgcd-/-; Mapk3-/-; Mapk1f/f
Sgcd-/-; Mapk3-/-; Mapk1f/f-Pax7Cre-ER
C
Central nuclei (%)
80
60
40
20
0
#
Quad.
Sgcd-/-; Mapk3-/-; Mapk1f/f
Sgcd-/-; Mapk3-/-; Mapk1f/f-Pax7Cre-ER
D
IgM+ fibers/area
0.16
0.12
0.08
0.04
0
#
Quad.
E
Fibers (%)
50
40
30
20
10
0
Quad.
0-20
21-30
31-40
41-60
61-150
Feret's diameter (µm)

**Supplemental Figure 3**. Sustained protection from muscle damage in *Sgcd*$^{-/-}$ mice lacking satellite cells. (A) Schematic representation of the tmx treatment regimen. Two-month-old mice received a daily tmx injection for 3 consecutive days and were subsequently placed on tmx chow after weaning. Samples were collected at 7 months of age. (B) Representative H&E stained histological sections of the quadriceps from 7-month-old mice of the indicated genotypes. Scale bar = 100 µm. (C) Quantification of the number of myofibers with centrally located nuclei in the quad muscle from mice of the indicated genotypes. n = 4 for both groups. Significance was determined using a Student's t-test, # $P < 0.05$. (D) Quantification of IgM positive fibers in muscle sections from mice of the indicated genotypes. n = 3, *Sgcd*$^{-/-}$; *Mapk3*$^{-/-}$; *Mapk1*$^{f/f}$; n = 6, *Sgcd*$^{-/-}$; *Mapk3*$^{-/-}$; *Mapk1*$^{f/f\text{-}Pax7Cre\text{-}ER}$. Significance was determined using a Student's t-test, #$P < 0.05$. (E) Minimal feret's diameter distribution from the quad muscle of 7-month-old mice with the indicated genotypes. n = 5 for all groups. Data represent mean ± SEM for all graphs.

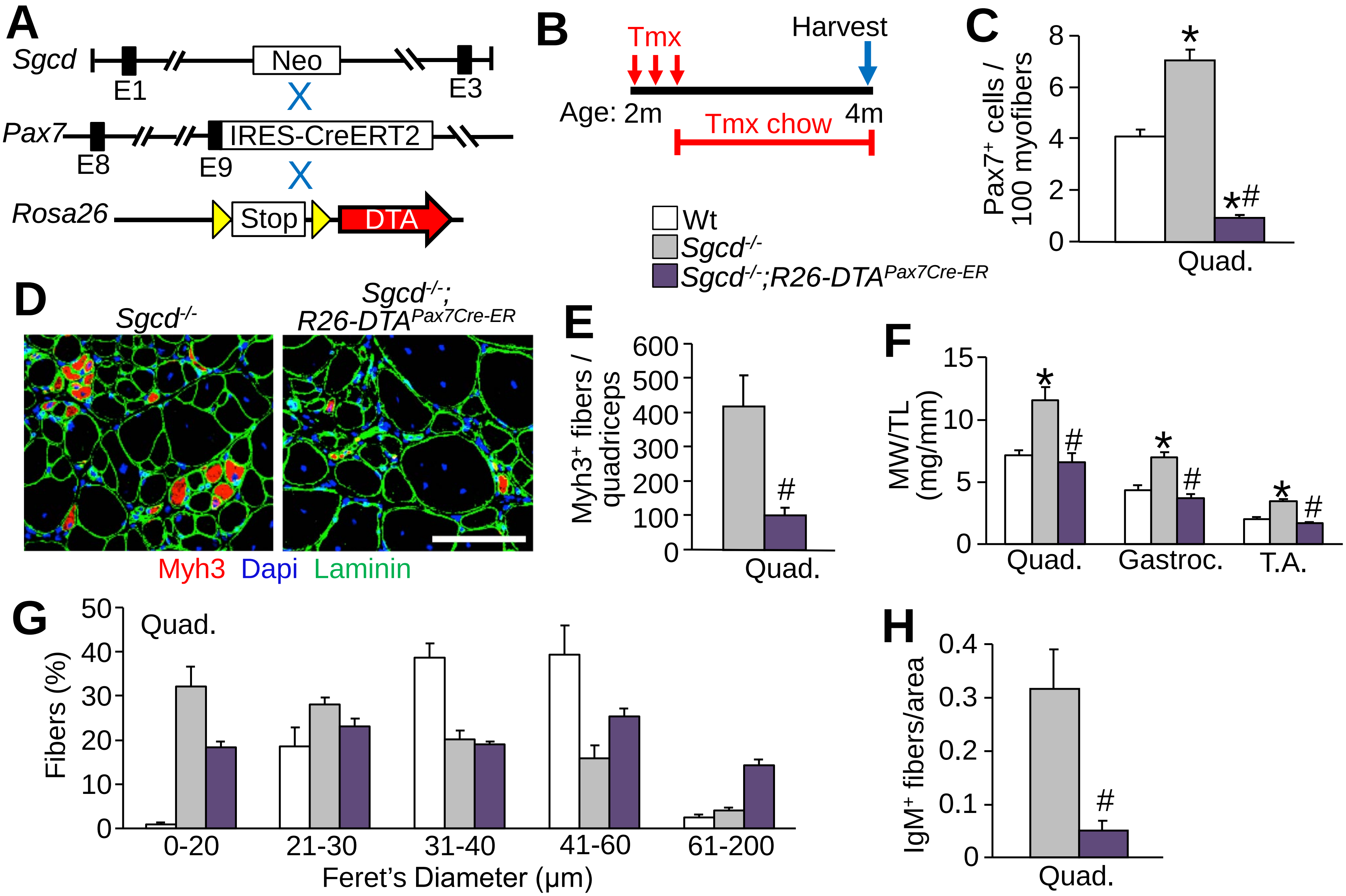
A
Sgcd
E1
Neo
E3
Pax7
E8
E9
IRES-CreERT2
Rosa26
Stop
DTA
B
Tmx
Harvest
Age: 2m
4m
Tmx chow
C
Pax7+ cells / 100 myofibers
Quad.
Wt
Sgcd-/-
Sgcd-/-;R26-DTAPax7Cre-ER
D
Sgcd-/-
Sgcd-/-; R26-DTAPax7Cre-ER
Myh3 Dapi Laminin
E
Myh3+ fibers / quadriceps
Quad.
F
MW/TL (mg/mm)
Quad.
Gastroc.
T.A.
G
Quad.
Fibers (%)
0-20
21-30
31-40
41-60
61-200
Feret's Diameter (μm)
H
IgM+ fibers/area
Quad.

**Supplemental Figure 4.** Satellite cell ablation with DTA in *Sgcd*$^{-/-}$ mice. (A) Schematic of breeding the tmx-inducible *Pax7*$^{Cre\text{-}ERT2}$ mice with *Rosa26-DTA* targeted mice. These lines were crossed onto the *δ-sarcoglycan*–null (*Sgcd*$^{–/–}$) background. (B) Schematic representation of the tmx treatment regimen. Two-month-old mice received a daily tmx injection for 5 consecutive days and were subsequently placed on tmx chow until harvest. (C) Quantification of Pax7 positive satellite cells in muscle sections from the quad of mice with the indicated genotypes. n = 4 for all groups. A one-way ANOVA with Tukey's multiple comparisons test was used to determine significance, *P < 0.05 versus wt, # P < 0.05 versus disease controls. The 4-month wt and *Sgcd*$^{-/-}$ Pax7 positive counts are also shown in Figure 4. (D) Representative quad muscle sections immunostained for Myh3 (red) and laminin (green) in 4-month-old mice of the indicated genotypes. Dapi stained nuclei are in blue. Scale bar = 100 μm. (E) Quantification of the number of Myh3 positive fibers in quad muscle sections of mice with the indicated genotypes. n = 4 for both groups. Significance was determined using a Student's t-test, #P < 0.05. The 4-month *Sgcd*$^{-/-}$ Myh3 positive fiber counts are also shown in Figure 4. (F) M.W./T.L. ratios from mice of the indicated genotypes at 4 months of age. n = 8, wt; n = 4, *Sgcd*$^{-/-}$; n = 4, *Sgcd*$^{-/-}$; *R26-DTA*$^{Pax7Cre\text{-}ER}$. A one-way ANOVA with Tukey's multiple comparisons test was used to determine significance, *P < 0.05 versus wt, # P < 0.05 versus *Sgcd*$^{-/-}$. The 4-month wt and *Sgcd*$^{-/-}$ weight data are also shown in Figure 4. (G) Minimal feret's diameter distribution from the quad muscle of 4-month-old mice with the indicated genotypes. The 4-month wt and *Sgcd*$^{-/-}$ feret's diameter data are also shown in Figure 4. (H) Quantification of IgM positive fibers in muscle section from mice of the indicated genotypes. n = 5, *Sgcd*$^{-/-}$; n = 4, *Sgcd*$^{-/-}$; *R26-DTA*$^{Pax7Cre\text{-}ER}$. Significance was determined using a t-test, #P < 0.05. Data represent mean ± SEM for all graphs.

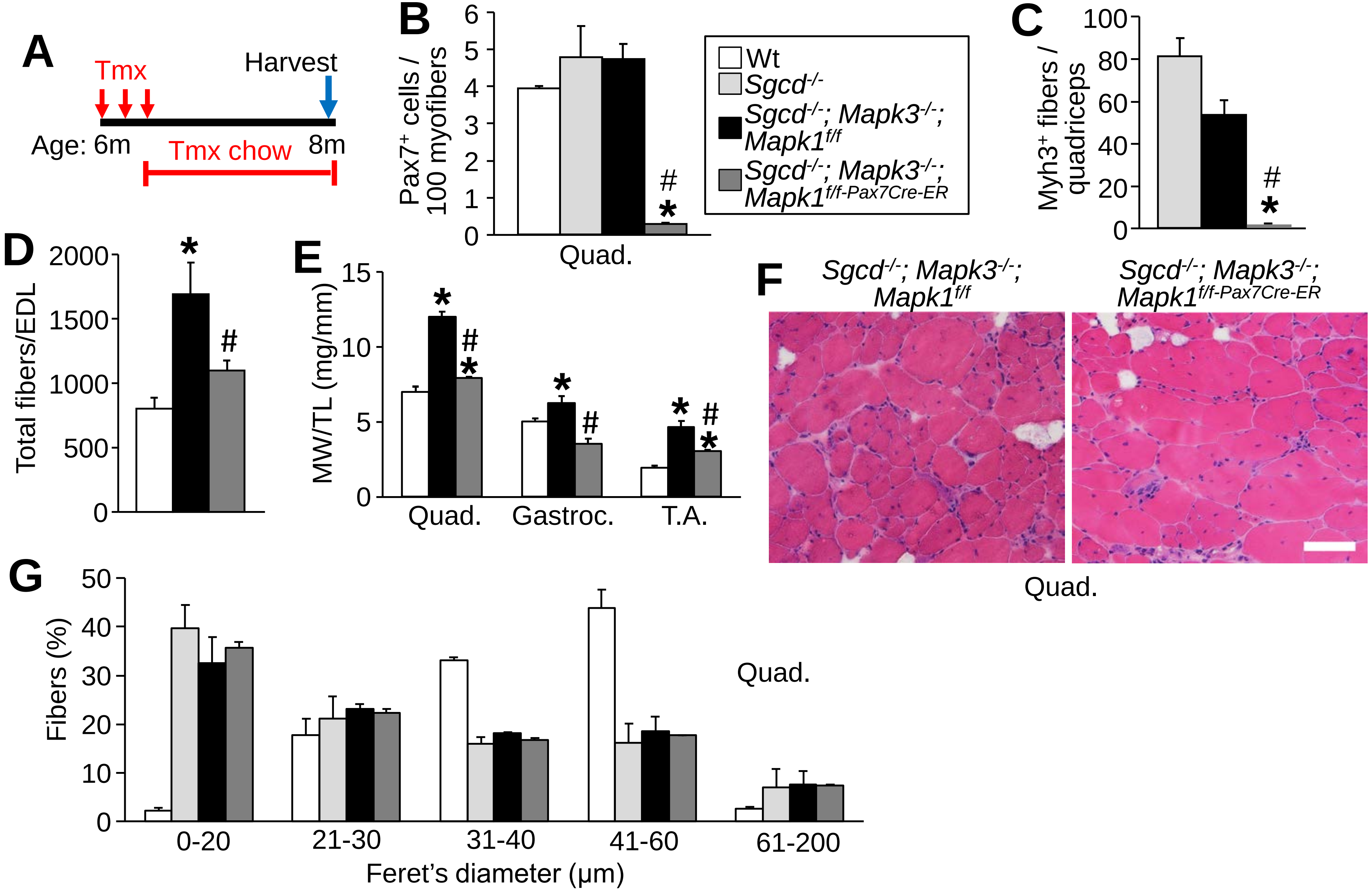
A
Tmx
Harvest
Age: 6m
Tmx chow
8m
B
Pax7+ cells / 100 myofibers
Quad.
Wt
Sgcd-/-
Sgcd-/-; Mapk3-/-; Mapk1f/f
Sgcd-/-; Mapk3-/-; Mapk1f/f-Pax7Cre-ER
C
Myh3+ fibers / quadriceps
D
Total fibers/EDL
E
MW/TL (mg/mm)
Quad.
Gastroc.
T.A.
F
Sgcd-/-; Mapk3-/-; Mapk1f/f
Sgcd-/-; Mapk3-/-; Mapk1f/f-Pax7Cre-ER
Quad.
G
Fibers (%)
Quad.
0-20
21-30
31-40
41-60
61-200
Feret's diameter (µm)

**Supplemental Figure 5**. Satellite cell ablation in adult *Sgcd*$^{-/-}$ mice. (A) Schematic representation of the tamoxifen (tmx) treatment regimen. Adult (6-month-old) mice received a daily tmx injection for 3 consecutive days and were subsequently placed on tmx chow. (B) Quantification of satellite cell numbers from quadriceps (quad.) muscle sections from 8-month-old mice of the indicated genotypes. n = 3, wt; n = 4 *Sgcd*$^{-/-}$; n = 3, *Sgcd*$^{-/-}$; *Mapk3*$^{-/-}$; *Mapk1*$^{f/f}$; n = 3, *Sgcd*$^{-/-}$; *Mapk3*$^{-/-}$; *Mapk1*$^{f/f\text{-}Pax7Cre\text{-}ER}$. A one-way ANOVA with Tukey's multiple comparisons test was used to determine significance, *P < 0.05 versus wt, # P < 0.05 versus disease controls. (C) Quantification of the number of Myh3 positive fibers in quad muscle sections of mice with the indicated genotypes. n = 3, wt; n = 4 *Sgcd*$^{-/-}$; n = 3, *Sgcd*$^{-/-}$; *Mapk3*$^{-/-}$; *Mapk1*$^{f/f}$; n = 3, *Sgcd*$^{-/-}$; *Mapk3*$^{-/-}$; *Mapk1*$^{f/f\text{-}Pax7Cre\text{-}ER}$. A one-way ANOVA with Tukey's multiple comparisons test was used to determine significance, # P < 0.05 versus disease controls. (D) Quantification of the total myofibers present in the extensor digitorum longus (EDL) muscle from 8-month-old mice of the indicated genotypes. n = 6, wt; n = 3, *Sgcd*$^{-/-}$; *Mapk3*$^{-/-}$; *Mapk1*$^{f/f}$; n = 3, *Sgcd*$^{-/-}$; *Mapk3*$^{-/-}$; *Mapk1*$^{f/f\text{-}Pax7Cre\text{-}ER}$. A one-way ANOVA with Tukey's multiple comparisons test was used to determine significance, *P < 0.05 versus wt, # P < 0.05 versus *Sgcd*$^{-/-}$; *Mapk3*$^{-/-}$; *Mapk1*$^{f/f}$. (E) M.W./T.L. ratios from mice of the indicated genotypes at 8 months of age. n = 6, wt; n = 3, *Sgcd*$^{-/-}$; *Mapk3*$^{-/-}$; *Mapk1*$^{f/f}$; n = 3, *Sgcd*$^{-/-}$; *Mapk3*$^{-/-}$; *Mapk1*$^{f/f\text{-}Pax7Cre\text{-}ER}$ A one-way ANOVA with Tukey's multiple comparisons test was used to determine significance, *P < 0.05 versus wt, # P < 0.05 versus *Sgcd*$^{-/-}$; *Mapk3*$^{-/-}$; *Mapk1*$^{f/f}$. (F) Representative H&E stained histological sections of the quad muscle from mice of the indicated genotypes. Scale bars = 100 μm. (G) Minimal feret's diameter distribution from the quad muscle of 8-month-old mice with the indicated genotypes. n = 3 for all groups. Data represent mean ± SEM for all graphs.
